# Functional decoding reveals a hidden regulatory layer of the *Salmonella* transcriptome during infection

**DOI:** 10.64898/2026.09.02.748214

**Authors:** Goloventzitz Izchak, Maya Elgrably-Weiss, Fayyaz Hussain, Jens Georg, Shoshy Altuvia

## Abstract

Bacterial transcriptomes contain extensive, largely unexplored regulatory information beyond annotated genes, including small regulatory RNAs (sRNAs), yet distinguishing functionally active transcripts from the broader non-coding transcriptome remains a fundamental challenge, particularly in the context of host-pathogen interactions. Here, we develop an unbiased high-throughput functional screening strategy to decode the regulatory potential of the *Salmonella* enterica transcriptome during macrophage infection. A pooled expression library comprising 875 RNA fragments derived from infection-relevant conditions was screened for effects on bacterial invasion and intracellular survival, revealing distinct and stage-specific regulatory activities. Functional characterization uncovered non-canonical sRNAs originating from 5ʹ untranslated regions that differentially modulate virulence-associated programs spanning the SPI-1, SPI-4 and SPI-2 pathogenicity islands. One attenuates the SPI-1 and SPI-4 secretion systems, limiting bacterial adhesion and invasion, whereas the other promotes invasion-associated programs while repressing pathways supporting intracellular survival. Rather than acting solely on individual virulence determinants, these regulatory activities reveal a broader layer of post-transcriptional control that coordinates bacterial adaptation across the dynamic host environment. Our study establishes a scalable strategy for extracting functional regulatory information from condition-specific bacterial transcriptomes and provides a framework for uncovering context-dependent RNA-mediated control of complex host-associated phenotypes.

## Introduction

The ability of bacteria to adapt to rapidly changing environments depends on their capacity to dynamically interpret and remodel gene expression. While protein-coding genes constitute the principal focus of bacterial genome annotation, increasing evidence indicates that substantial regulatory information resides outside annotated coding sequences, within non-coding transcripts and untranslated regions. Small regulatory RNAs (sRNAs) represent one of the best-characterized classes of such regulatory elements, acting post-transcriptionally to modulate the translation and stability of target mRNAs. Over the past three decades, large-scale transcriptomics, comparative genomics, and genetic screens have identified hundreds of sRNAs across diverse bacterial species, establishing them as central regulators of bacterial physiology, stress responses, and virulence (Holmqvist & Wagner, Gottesman & Storz, Papenfort & Vogel) (Papenfort & Melamed, Vogt & Fröhlich, Papenfort & Storz, Schnoor *et al*., Banerjee).

As bacterial transcriptomes have been explored in increasing depth, numerous RNAs have been identified that originate from unexpected genomic contexts, including protein-coding genes, untranslated regions (UTRs), and internal coding sequences. (Krieger *et al*., Adams *et al*.). Many of these RNAs are generated through premature transcription termination, RNA processing, or internal promoter activity (Chao & Vogel, Hör *et al*., Ponath *et al*.). Complementary RNA-seq and global RNA interactome studies have revealed that such fragments are abundant and frequently stable (Adams *et al*., Chao *et al*., Kröger *et al*., Guo *et al*., Holmqvist *et al*., Updegrove *et al*., Wang *et al*., Hoyos *et al*., Liu *et al.,* Melamed *et al*.). Yet transcript abundance alone does not establish regulatory function. Consequently, distinguishing functional regulatory RNAs from pervasive transcription has emerged as a major challenge, particularly in the context of host infection.

Addressing this challenge requires functional rather than descriptive approaches. However, conventional overexpression, deletion, and mutational analyses rely on one-by-one characterization of individual candidates, making them labor-intensive and poorly suited for systematically identifying functional sRNAs and defining their biological roles during complex environments including host-pathogen interactions (Barquist & Vogel, Saliba *et al*.). As a result, despite rapidly expanding catalogs of bacterial sRNAs, functional discovery has remained largely candidate-driven.

*Salmonella enterica,* a facultative intracellular pathogen, provides an ideal model to address this problem. More than 280 sRNAs are expressed under infection-relevant conditions (Kröger *et al*.), yet only a fraction have been functionally linked to virulence (Sittka *et al*., Padalon-Brauch *et al*., Gong *et al*., Ryan *et al*., Wang *et al*., Westermann *et al*., Kim *et al*., Kim *et al*., Abdulla *et al*.).

Successful infection depends on the sequential activation of pathogenicity island (SPI)-encoded virulence programs that mediate epithelial invasion, intracellular survival, and dissemination (Ilyas *et al*., Li *et al*.). Although the transcriptional regulation of these programs has been extensively characterized, whether additional sRNAs coordinate these stage-specific transitions remains largely unknown.

To overcome the limitations of conventional sRNA discovery, we developed a scalable, unbiased, phenotype-based strategy for systematically identifying functionally active sRNAs. We generated a plasmid library expressing short RNA fragments derived from *Salmonella* grown under infection-mimicking conditions and screened this library during macrophage infection. This unbiased approach revealed 51 RNA elements that impair bacterial invasion and/or intracellular survival, uncovering a previously unexplored layer of infection-associated regulation. We subsequently characterized two previously unannotated 5ʹ UTR-derived sRNAs, EsvA and EsvB, which differentially regulate SPI-dependent virulence programs and coordinate bacterial functions across distinct stages of infection.

Current approaches to sRNA discovery typically rely on genomic annotation, sequence conservation, expression profiles, or predicted RNA features, making functional discovery dependent on prior assumptions about transcript identity and regulatory potential. Directly linking transcript-derived sequences to phenotypic function without such constraints could provide a complementary strategy for uncovering regulatory elements embedded within the bacterial transcriptome. Here, we establish a scalable, phenotype-driven framework for unbiased functional discovery of bacterial sRNAs directly from complex infection phenotypes. By extending discovery beyond canonical intergenic regulators, our approach reveals previously inaccessible regulatory elements and provides a general framework for dissecting RNA-mediated regulatory networks in bacterial pathogens.

## Results

### A comprehensive expression library of infection-enriched *Salmonella* RNAs enables functional screening

Most functional screening approaches in bacteria rely on genome annotation, making them poorly suited for the systematic discovery of regulatory RNAs, particularly non-canonical sRNAs that arise from untranslated regions, coding sequences, or RNA processing events. To overcome this limitation, we developed an annotation-independent, transcriptome-derived gain-of-function strategy. We generated a high-complexity plasmid expression library from short RNA fragments (100-400 nt) derived from the *Salmonella* enterica transcriptome under infection-mimicking conditions, enabling unbiased functional screening of expressed RNAs irrespective of their genomic origin or prior annotation. RNA fragments were cloned between the inducible PlacO-1 promoter and a strong transcriptional terminator, and the resulting plasmid library was transformed into a LacI-expressing *Salmonella* strain. Next-generation sequencing (NGS) followed by mapping revealed 875 unique sRNA candidates spanning diverse genomic contexts: ∼25% mapped downstream of annotated promoters, ∼7% mapped near stop codons, and ∼18% mapped as sense fragments within coding sequences, including some overlapping adjacent genes. Antisense fragments to 5ʹ and 3ʹ regions accounted for ∼8% and ∼7%, respectively. A smaller fraction mapped to intergenic regions or known sRNAs. (Figure 1A).

**Figure 1.**
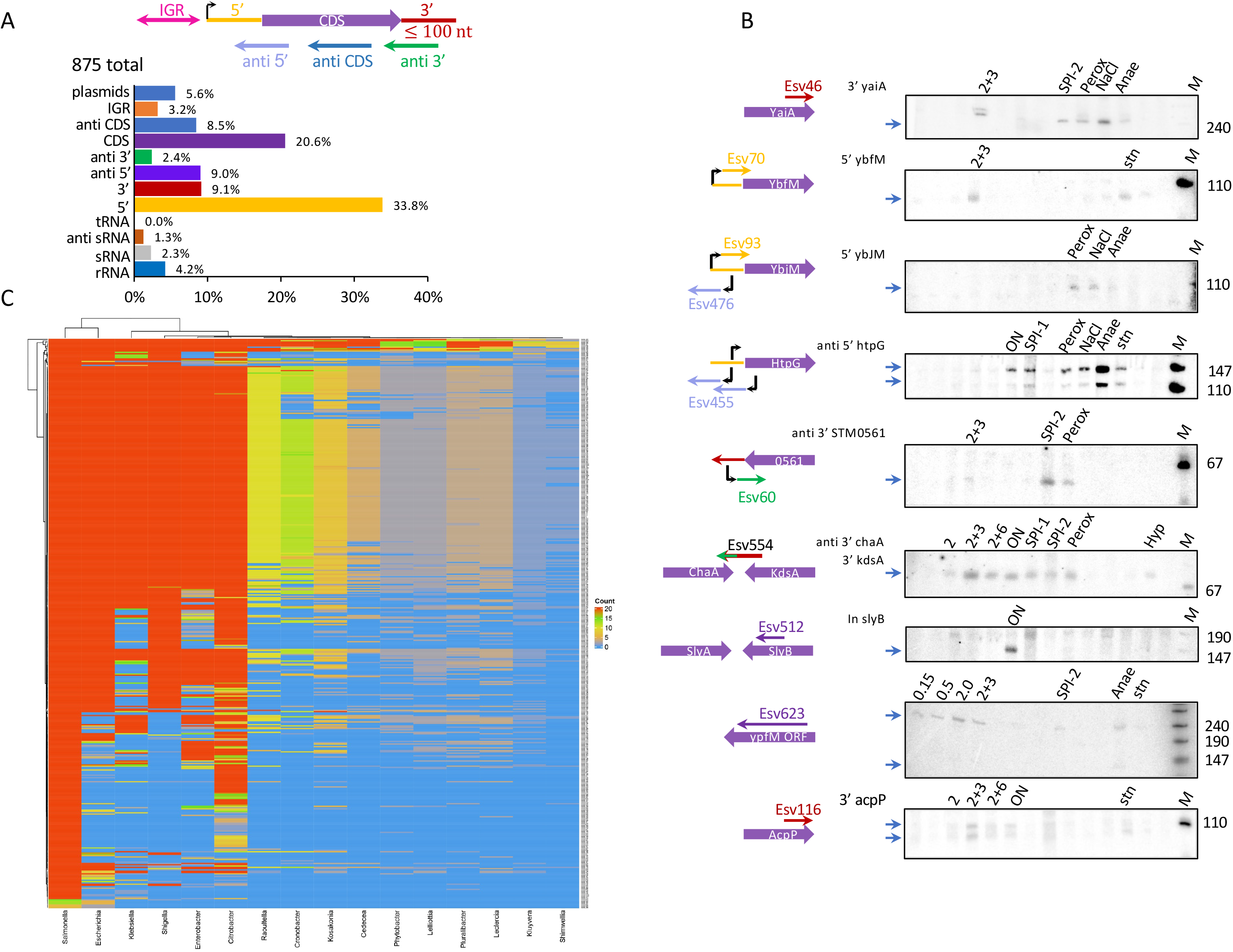
A comprehensive *Salmonella* sRNA expression library comprising 875 unique sRNA candidates derived from diverse genomic contexts. (A) Schematic genome-wide distribution of chromosomally encoded sRNA fragments cloned under infection-permissive conditions. (B) Northern blot validation of sRNA expression under defined conditions; tmRNA served as loading control (Figure S2). The colors correspond to the genomic distribution shown in panel A. (C) Conservation analysis (minimum column sum ≥40) revealed that 439 fragments are conserved (65%) across enterobacterial genera. 450 fragments (minimum column sum ≥1000) are conserved (57%) within *Salmonella* subspecies (Figure 1C, S3), highlighting their potential functional relevance. Interactive Shiny-based maps were used for visualization.

To confirm that the library represents the native *in vivo* landscape of active small RNAs, library fragments were compared to existing *Salmonella* transcriptomic data (Kröger *et al*.). Many fragments corresponded with previously detected RNA species, supporting their validity (Figure S1). Notably, 142 previously unannotated fragments were mapped within 10 nucleotides of known transcription start sites, and 94 fragments were located in proximity to RNase E cleavage sites (Chao *et al*.), further supporting the likelihood of their genuine existence. Northern blots confirmed the expression of multiple library fragments under various growth and stress conditions (Figure 1B, S2). The presence of many fragments derived from *Salmonella* grown under infection-mimicking (“standing”) conditions across multiple additional growth conditions (Kröger *et al*.) highlights the richness and diversity of the library, enabling functional studies across a wide range of environments. Importantly, by design, the library captures qualitative diversity rather than merely quantitative variation.

Conservation analysis revealed that 439 fragments exhibit greater than 65% sequence conservation across the enterobacteria genera, while 654 exhibit 57% conservation among *Salmonella* subspecies (Figure 1C, S3), underscoring potential functional importance (Table S1).

### Functional screening in macrophages identifies infection-stage-specific regulatory RNAs

*Salmonella* harbors multiple pathogenicity islands (SPIs) that are crucial for its survival and virulence, facilitating colonization, invasion, and persistence in hostile environments (Ilyas *et al*., Han *et al*., Sia *et al*.). Aligning the short RNAs from the library to the *Salmonella* genome, we observed a rise in the relative quantity of sRNAs located within certain pathogenicity islands, including SPI-1, SPI-2, SPI-5, and SPI-14, which regulate invasion and intracellular survival (Table 1). The enrichment was determined considering the size of the islands, with a cutoff ratio exceeding 1.5.

**Table 1.** Genomic islands enriched for sRNAs.

| Island | Size<sup>a</sup> | No. of sRNAs<sup>b</sup> | Ratio<sup>c</sup> | Library ratio<sup>d</sup> | Enrichment<sup>e</sup> |
| --- | --- | --- | --- | --- | --- |
| SPI-1 | 42204 | 15 | 4E-04 | 1.5E-04 | **2.4** |
| SPI-2 | 40165 | 10 | 2E-04 | 1.5E-04 | **1.7** |
| SPI-3 | 17022 | 2 | 1E-04 | 1.5E-04 | 0.8 |
| SPI-4 | 24750 | 4 | 2E-04 | 1.5E-04 | 1.1 |
| SPI-5 | 9109 | 7 | 8E-04 | 1.5E-04 | **5.1** |
| SPI-6 | 46314 | 7 | 2E-04 | 1.5E-04 | 1.0 |
| SPI-9 | 16681 | 0 | 0E+00 | 1.5E-04 | 0.0 |
| SPI-11 | 8413 | 2 | 2E-04 | 1.5E-04 | 1.6 |
| SPI-12 | 5266 | 0 | 0E+00 | 1.5E-04 | 0.0 |
| partial SPI-13 | 7478 | 0 | 0E+00 | 1.5E-04 | 0.0 |
| SPI-14 | 7372 | 2 | 3E-04 | 1.5E-04 | **1.8** |
| SPI-16 | 4198 | 0 | 0E+00 | 1.5E-04 | 0.0 |
| SLP105 | 45242 | 3 | 7E-05 | 1.5E-04 | 0.4 |
| bacteriophage SLP203 | 40088 | 5 | 1E-04 | 1.5E-04 | 0.8 |
| Oaf O-antigen modification locus | 9905 | 1 | 1E-04 | 1.5E-04 | 0.7 |
| CS54 genomic island | 24139 | 2 | 8E-05 | 1.5E-04 | 0.6 |
| SLP272 | 50513 | 11 | 2E-04 | 1.5E-04 | **1.5** |
| degenerate bacteriophage SLP281 | 10534 | 0 | 0E+00 | 1.5E-04 | 0.0 |
| SLP285 | 32907 | 6 | 2E-04 | 1.5E-04 | 1.2 |
| prophage SLP289 | 11907 | 3 | 3E-04 | 1.5E-04 | **1.7** |
| prophage remnant SLP443 | 22114 | 2 | 9E-05 | 1.5E-04 | 0.6 |

<sup>a</sup>Size of the island in bases<sup>b</sup>Number of short RNA fragments identified within an island<sup>c</sup>Ratio of short RNA fragments to island size<sup>d</sup>Ratio of total short RNA fragments to total genome: 732/4878013.<sup>e</sup>Enrichment - ratio of c to d
rRNA and tRNA were excluded from these analyses.
Areas with enrichment higher than 1.5 are highlighted in red.

Given the enrichment of SPI-associated transcripts in the library, we reasoned that it would be particularly well suited for functional screening under infection conditions. To identify functional sRNAs affecting infection, RAW 264.7 macrophages were infected with *Salmonella* expressing the library. Expression of the library fragments was induced prior to infection and maintained throughout. Bacteria were collected at 0, 45 min, and 8 h post-infection for NGS analysis (Figure 2A).

**Figure 2.**
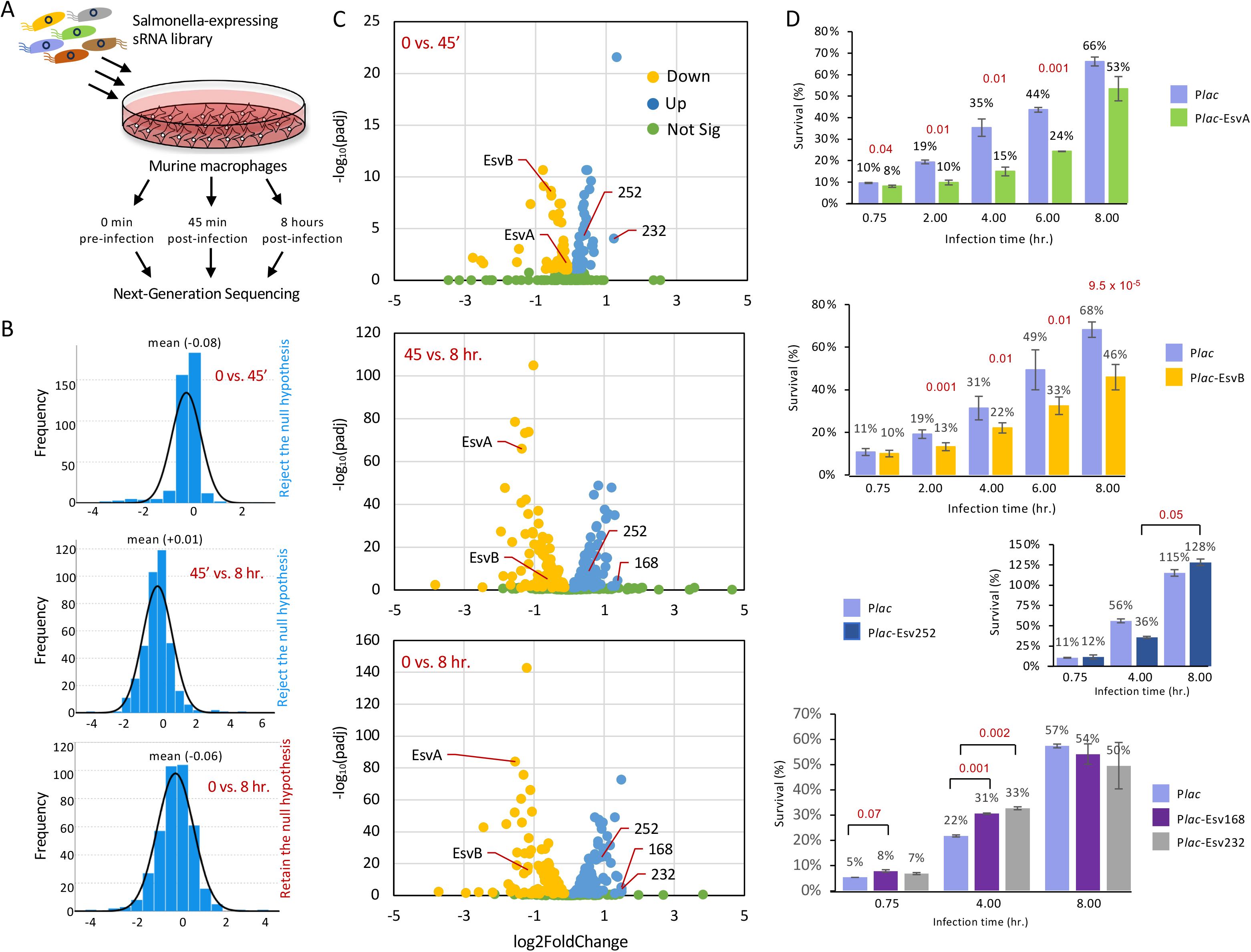
Functional screening in macrophages identifies infection-stage-specific regulatory RNAs (A) murine RAW 264.7 macrophages were infected with *Salmonella* expressing the sRNA plasmid library as described in Material and Methods. Expression of the sRNA library was induced prior to infection and maintained throughout. Bacteria were collected at 0, 45 min, and 8 h post-infection for NGS analysis. (B) Gaussian distribution analyses revealed a negative skew during invasion phase (0-45 min), consistent with sRNAs that limit entry, and a positive skew during the intracellular survival phase (45 min-8 h). (C) Volcano plot analysis of 395 fragments present at the selective infection identifies 38 fragments that enhanced survival and 51 that impaired it (P ≤ 0.1; fold-change ≥ 1.6). Only a subset of these fragments is highlighted. (D) Macrophage survival assays confirmed that EsvA and EsvB reduced *Salmonella* survival, whereas Esv252 promoted late intracellular survival and Esv168 and Esv232 sRNAs enhanced both entry and intracellular survival, both competitively and non-competitively. Data represent mean ± SD of 3-6 biological replicates; *p*-values (two-tailed unpaired t-test) are indicated.

Infection of RAW 264.7 macrophages with the sRNA library revealed dynamic shifts in fragment abundance across distinct stages of infection. Gaussian distribution analyses showed a negative skew during the early phase of infection (0-45 min), consistent with enrichment of sRNAs that may restrict host-cell entry, followed by a positive skew during the later phase (45 min to 8 h), suggesting the involvement of sRNAs that may influence intracellular survival (Figure 2B)

Differential abundance analysis showed that, among of the 395 fragments present during selective infection (0 vs. 8 hours), 38 enhanced survival, whereas 51 impaired it (padj ≤ 0.1; fold-change ≥ 1.6) (Figure 2C). We prioritized two 5ʹ UTR-derived sRNAs for mechanistic dissection: EsvA and EsvB named for *Experimentally discovered Small RNAs associated with Virulence*, which regulate genes associated with invasion and/or intracellular survival (Figure 2D).

While the EsvA and EsvB sRNAs were associated with reduced *Salmonella* survival in macrophages, Esv168 and Esv232 enhanced host cell entry and intracellular survival, whereas Esv252 promoted intracellular survival at later stages of infection, both through competitive (in a library pool) and non-competitive mechanisms (as a single strain infection) (Figure 2CD).

### EsvA represses SPI-4, an adhesion-associated locus

The 160-nucleotide long EsvA is derived from the 5’ region of STM0327, consisting of 35 nucleotides of STM0327 untranslated region (UTR) and 125 nucleotides of the coding sequence (CDS) (Figure 3A). STM0327 encodes a conserved hypothetical protein essential for survival in macrophages (Chan *et al*.). The chromosomally encoded short RNA fragment EsvA is detectable during stationary phase (OD 600 = 3.0), under-standing growth and hypoxic conditions, (Figure 3B). Its stability is influenced by RNase III: in *rnc* mutants, both the full-length species and a processed form may accumulate, whereas in some cases only the processed form is visible. (Figure 3B, S4B). In contrast, RNase E does not appear to have any effect on the generation of EsvA. (Figure S4B). In addition, we found that plasmid-encoded *in trans* expression of sRNA EsvA increases the transcript levels of the chromosomally encoded STM0327 (Figure S4C). Rifampicin treatment showed that EsvA RNA is highly unstable (Figure S4D, lanes 6-8). Notably, its presence influences the stability of STM0327 mRNA (Figure S4D, lanes 7,8). In the absence of EsvA, STM0327 mRNA levels are markedly reduced, suggesting that EsvA expressed in trans may sequester a factor involved in RNA processing or degradation.

**Figure 3.**
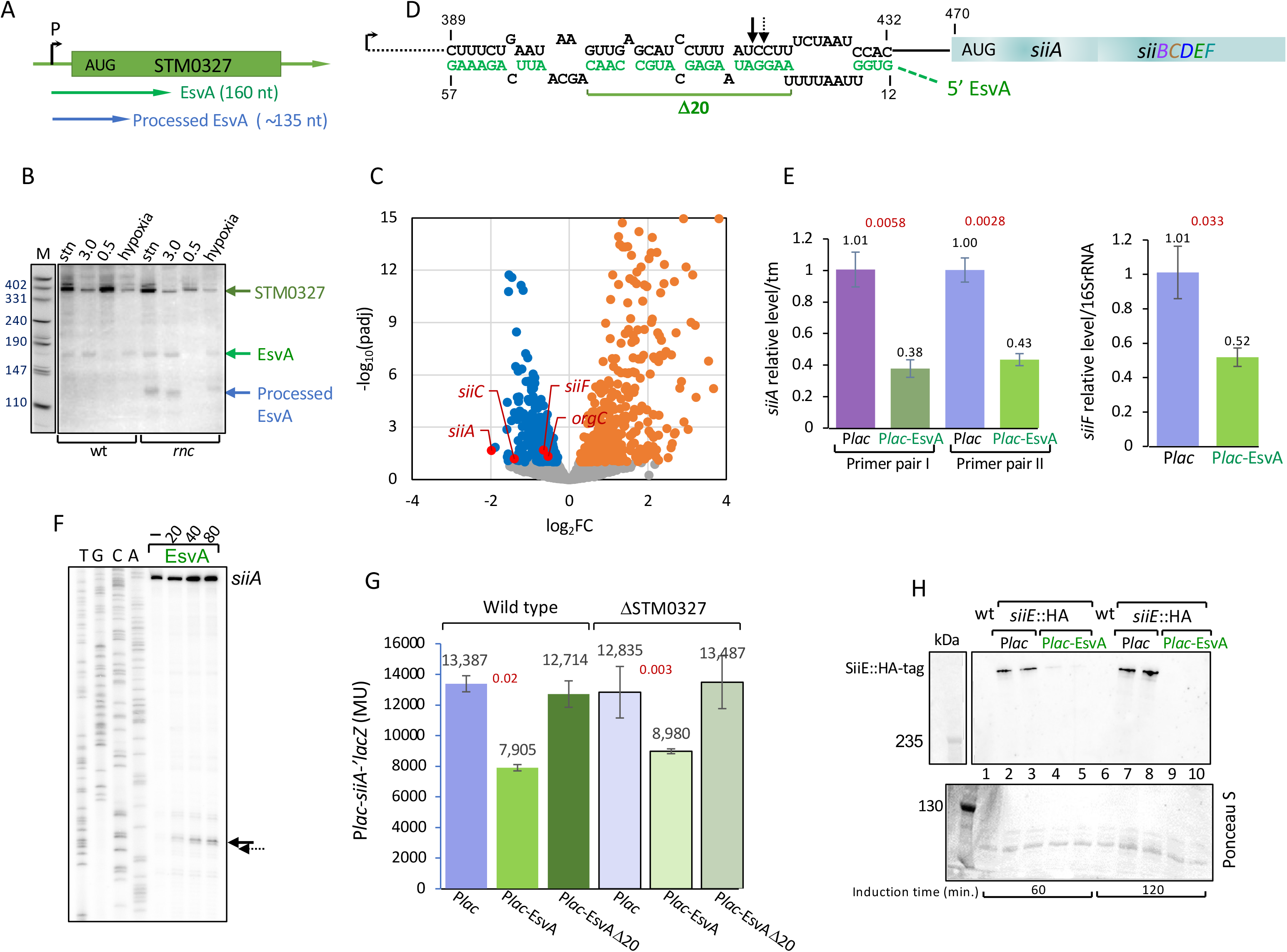
EsvA originates from the 5’UTR of STM0327 (A) Schematic representation of EsvA and STM0327. The green arrow indicates the position and orientation of EsvA. (B) The chromosomally encoded short RNA fragment EsvA is detectable under the indicated conditions. Its stability is positively influenced by RNase III (*rnc*) (Figure S4B). (C) Pulse expression of plasmid-encoded EsvA in wild-type cells, followed by transcriptome analysis, identified *siiA* as a major target. The volcano plot highlights three genes of the *siiA-siiF* operon encoded within SPI-4. (D) Potential RNA-RNA interaction between the 5ʹ untranslated region (UTR) of *siiA* and EsvA predicted using the IntaRNA algorithm. The green bar indicates an internal 20-nt deletion in EsvA. (E) Quantitative real-time PCR analysis of *siiA* and *siiF*, representing the first and last genes of the operon. *siiA* was examined with two different sets of primers. Data represent mean ± SD of 3-6 biological replicates; *p*-values (two-tailed unpaired t-test) are indicated. (F) *In vitro* primer extension of *siiA* RNA in the presence of increasing concentrations of EsvA RNA revealed a major termination site within the predicted hybrid. (G) Translational *siiA-lacZ* fusion assays showed a decrease in reporter activity upon EsvA expression. A similar decrease in *siiA*-*lacZ* activity was detected in ΔSTM0327. EsvA variant carrying an internal 20-nucleotide deletion exhibited no detectable regulatory effect, confirming the functional importance of this interaction site. Data represent mean ± SD of 3-6 biological replicates; *p*-values (two-tailed unpaired t-test) are indicated. (H) Tagged SiiE::HA exhibits a decrease SiiE::HA protein levels due to in trans expression of EsvA. Western and Ponceau S staining as loading control.

In macrophages, EsvA overexpression is detrimental in cells carrying the STM0327 wild-type allele, mirroring its effect in STM0327-deficient cells. These findings suggest that EsvA regulates genes involved in macrophage infection, independently of STM0327. (Figure S4F).

Pulse expression of plasmid-encoded EsvA or STM0327 in wild-type cells, followed by transcriptome analysis (Figure S5) identified *siiA* and *siiF* as major targets (Figure 3C). *siiA*, the first gene of the SPI-4 *siiABCDEF* operon, encodes a type I secretion–associated protein that, together with SiiC, SiiD, and SiiF, forms the apparatus exporting SiiE, a _∼_ 600 kDa adhesin essential for invasion (Wagner *et al*.). Computational prediction of potential RNA-RNA interactions using the IntaRNA algorithm identified a putative binding site within the 5ʹ untranslated region (UTR) of *siiA*, located 38 nucleotides upstream of the translational start codon (Figure 3D). Quantitative real-time PCR analysis of *siiA* and *siiF*, the first and last genes of the operon, showed that EsvA expression reduced their transcript levels by 2.5-fold and 2-fold, respectively, indicating that EsvA modulates transcript abundance across the operon (Figure 3E).

To visualize the interaction between EsvA sRNA and *siiA* mRNA, in vitro primer extension assays were performed, which identified a single termination site within the predicted interaction region (Figure 3F). Supporting this interaction, translational *siiA-lacZ* fusion assays showed a corresponding decrease in reporter activity upon EsvA expression. Furthermore, an EsvA variant carrying an internal 20-nucleotide deletion (EsvAΔ20) showed no detectable regulatory activity, confirming the functional importance of this interaction site (Figure 3G). Notably, the EsvAΔ20 mutant also failed to stabilize STM0327 mRNA (Figure S4E), indicating that in trans expression of intact EsvA is required for STM0327 mRNA stabilization.

To assess the impact of EsvA on SiiE, an adhesin essential for invasion, we employed a tagged SiiE construct (SiiE::HA). Western blot analysis showed that EsvA expression markedly reduced SiiE abundance (Figure 3H and Figure S6). Collectively, these results indicate that EsvA downregulates the *siiA-siiF* operon encoded by SPI-4 to modulate *Salmonella* virulence. Consistently, expression of EsvA inhibits adhesion to polarized MDCK cells (Figure S7A), but not to unpolarized MDCK cells or to HeLa cells (Figure S7BC). This aligns with previous findings showing that the knockout of SPI-4 leads to impaired bacterial adhesion specifically to polarized epithelial cells ((Kiss *et al*., Galán & Curtiss, Gerlach *et al*.).

### EsvA represses SPI-1, an invasion associated locus

As *Salmonella* requires the cooperative activity of the SPI-4 encoded SiiE and the SPI-1 type III secretion system (T3SS) to mediate epithelial attachment and invasion (Kiss *et al*., Galán & Curtiss, Gerlach *et al*.), we postulated that additional genes affected by EsvA might influence SPI-1-dependent secretion or invasion. Indeed, while analyzing SiiE protein levels, we observed that the culture supernatant of cells expressing EsvA in trans was markedly different from that of the control (Figure 4A). Specifically, the supernatant from EsvA-expressing cells lacked both flagellar components and elements of the SPI-1 type III secretion system, indicating that EsvA impairs secretion of both the flagellar and SPI-1 type III secretion pathways. Consistent with these observations, both CopraRNA predictions (Table S2) and transcriptome analyses identified *orgB* and *orgBC* respectively, as candidate genes potentially regulated by EsvA. The *orgABC* locus is encoded within SPI-1, alongside *prgHIJK*. Moreover, individual mutations in *orgA* or *orgB* - the oxygen regulated invasion proteins, as well as in any of the *prg* genes, result in a general defect in *Salmonella* effector protein secretion and translocation (Klein *et al*., Sukhan *et al*.). Computational prediction of potential RNA-RNA interactions between the *orgABC* locus and EsvA using the IntaRNA algorithm identified two putative binding sites: one located within the *orgA* coding sequence (CDS) and another overlapping the *orgB* AUG start codon and the *orgA* stop codon (Figure 4B and S8AB). Quantitative real-time PCR analysis of *orgA*, *orgB*, and *orgC* revealed that EsvA expression reduced transcript levels by approximately 1.5-to 3-fold, indicating that EsvA modulates transcript abundance across this genetic locus (Figure 4 BC). To determine whether this regulatory effect extends to the protein level, we analyzed an OrgB-SPA fusion carrying a C-terminal sequential peptide affinity (SPA) tag (Zeghouf *et al*., 2004). OrgB–SPA levels were reduced upon EsvA expression (Figure 4D), consistent with the observed decrease in *orgB* transcript levels. Thus, EsvA impairs *Salmonella* adhesion and invasion of cells by repressing both the pathogenicity island 4 (SPI-4)-encoded Type I secretion system (T1SS) and the SPI-1-encoded Type III secretion system (T3SS).

**Figure 4.**
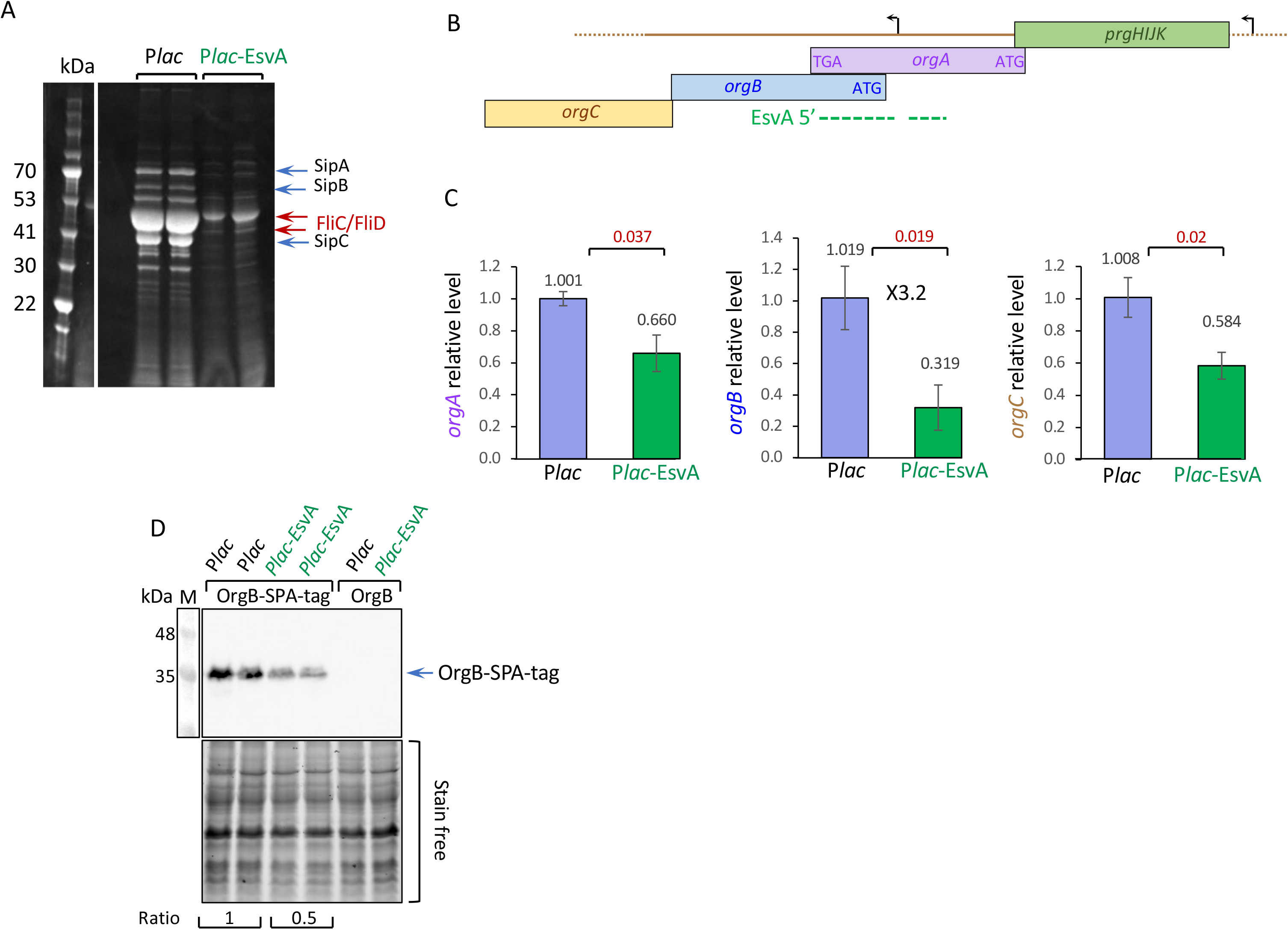
EsvA impairs SPI-1 type III secretion. (A) *In trans* expression of EsvA impairs secretion into the culture supernatant by both the flagellar and the SPI-1 type III secretion systems. Samples were taken 4 hours post EsvA induction and treated as described in the methods. (B) Schematic representation of the *prgHIJK-orgABC* locus, showing the position of the genes *orgA*, *orgB* and *orgC*. Arrows indicate promoter regions: one (far right) that drives transcription of the entire *prgHIJK-orgABC* operon, and another that specifically initiates transcription of *orgBC*. Dashed green line indicates predicted interactions with EsvA, see Figure S8 for details. (C) *In trans* ectopic expression of EsvA reduces mRNA levels of *orgA*, *orgB* and *orgC*. Quantitative real-time PCR analysis. Data represent mean ± SD of 3 biological replicates; p-values (two-tailed unpaired t-test) are indicated. (D) EsvA represses OrgB-SPA protein levels. Cultures of wild type and OrgB-SPA grown to OD600=0.3 were subjected to oxygen shock as described in the methods section. Wild type strains lacking SPA carrying P*lac* and P*lac*-EsvA served as controls.

### EsvB targets *pipB2*, an SPI-2 effector associated with intracellular survival, via *cis*-antisense interaction

EsvB is a 117-nucleotide sRNA derived from the 5ʹ untranslated region (UTR) of *pipB2* mRNA. *pipB2*, a type III secretion system effector encoded within *Salmonella* Pathogenicity Island 2 (SPI-2), contributes to *Salmonella*’s intracellular survival within macrophages (Henry *et al*., Han *et al*.). The *pipB2* transcript has been reported to contain three promoters-primary, secondary, and one located proximal to the *pipB2* start codon (P*pipB2*) (Kröger *et al*.). Our primer extension analysis confirmed the presence of three promoters and revealed a fourth start site at the 5ʹ end of EsvB, which may arise either from promoter activity or from RNA processing (Figures S9 ABCD). All promoters were active under high cell density and under conditions that induce SPI-2 expression (Figure S9 CD). Northern blot analysis revealed that, in addition to the short EsvB transcript, a longer transcript likely originating from the primary promoter and encompassing the EsvB fragment is also present. This longer RNA is particularly abundant at high cell density and at pH 5. In the *rnc* mutant, the short EsvB transcript was dramatically reduced, indicating that RNase III contributes to EsvB maturation, likely by cleaving a stem–loop structure in a transcript first processed by RNase E. (Figure S9E). Although the 5ʹ UTR of *pipB2* contains several RNase E cleavage sites, including sites located upstream of and proximal to the 5ʹ end of EsvB (Chao *et al*.) RNase E does not appear to be involved in generating the mature form of EsvB. Instead, EsvB is likely processed from a longer RNA transcript by RNase III. The appearance of even longer transcripts in addition to EsvB in the RNase G and RNase II mutants, suggest that these enzymes may contribute to transcript processing and stabilization (Figure S9E).

A CopraRNA (Wright *et al*.) search for EsvB targets identified 20 potential targets (false discovery rate ≤ 0.05), with *pipB2* emerging as the most significant. In addition, *sipB* and *ssaL*, two genes encoding components of two types of type III secretion system, were also ranked among the top targets (Table S3).

RNAfold analysis predicted that EsvB contains a long-inverted repeat, suggesting that one strand (positions 150-211) could act as an antisense and base-pair with the complementary strand (Figure 5BC). To test the interaction between EsvB and *pipB2*, we performed in vitro primer extension assays using two fragments representing the upper strand (Figure 5BC). The 5ʹ region of EsvB (positions 150-183, marked in red) bound *pipB2* RNA at two sites: one downstream of the start codon (DS) and another upstream of the AUG (US). By contrast, the adjacent sequence fragment showed no detectable binding to *pipB2* (Figure 5CDE). These results demonstrate that the *cis*-encoded EsvB acts as an antisense RNA that specifically targets *pipB2* mRNA. Quantitative real-time PCR analysis of *pipB2* carried out under high density and pH 5 conditions revealed that in-trans EsvB expression reduced transcript levels by approximately 2-fold and 5-fold, respectively suggesting that EsvB modulates *pipB2* transcript abundance (Figure 5F)

**Figure 5.**
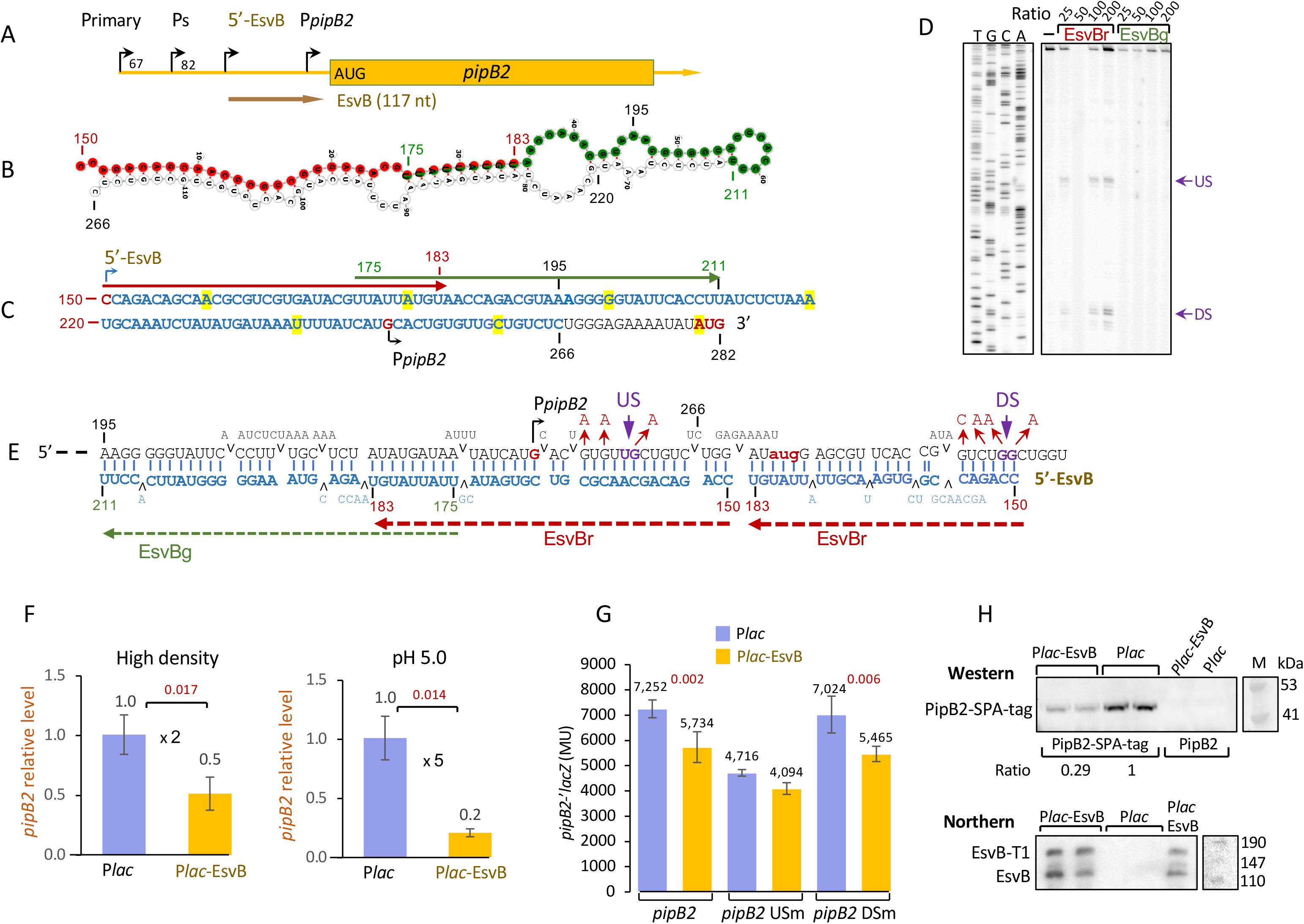
EsvB acts as an antisense RNA to inhibit *pipB2* expression. (A) Schematic representation of the *pipB2* transcript showing three previously identified promoters: primary (Primary), secondary (Ps), and one located just upstream of *pipB2* (P*pipB2*). Also indicated is the 5’ end of EsvB as determined by primer extension (see Figure S9C (B) Predicted secondary structure of EsvB showing a long-inverted repeat. (C) Structure prediction highlights two candidate binding regions at positions 150-183 (EsvBr-red) and 17-211 (EsvBg-green). (D-E) *In vitro* primer extension assays revealed that the 5ʹ EsvB fragment (150-183) binds *pipB2* at two sites, upstream (US) and downstream (DS) of the start codon. The hybrid structure is shown with EsvB (red dashed lines) pairing at US and DS (purple); red arrows mark mutations introduced to disrupt binding. (F) Quantitative real-time PCR analysis of *pipB2* carried out under high density and pH 5 conditions. *In trans* EsvB expression reduced transcript levels by approximately 2-fold and 5-fold, respectively. Data represent mean ± SD of 3 biological replicates; *p*-values (two-tailed unpaired t-test) are indicated. (G) Translational *pipB2-lacZ* fusions (wild type, US mutant, DS mutant) were tested for repression by *in trans* EsvB. EsvB repressed wild type and DS mutant, but not the US mutant, indicating that the upstream site is critical. Data represent mean ± SD of 5 biological replicates; *p*-values (two-tailed unpaired t-test) are indicated. (H) Effect of EsvB on PipB2 protein levels. A strain carrying C-terminal PipB2-SPA fusion (Zeghouf *et al*.) was analyzed by western blot. Cultures (OD600 of 0.2) were induced (1 mM IPTG, 120 min) to express EsvB. P*lac* and wild type *pipB2* served as controls. Northern blot confirmed EsvB expression (primer 4027); the upper band corresponds to readthrough transcription ending at T1.

To assess the post-transcriptional effect of EsvB on *pipB2*, we constructed a *pipB2*-*lacZ* translational fusion. Expression of EsvB in trans led to a reduction in *pipB2*-*lacZ* levels (Figure 5G). Mutation of the downstream (DS) site did not abolish repression, whereas mutation of the upstream (US) site impaired EsvB-mediated repression, indicating that the upstream site is critical for regulation (Figure 5G). The effect of EsvB on PipB2 protein levels was further validated using a PipB2-SPA fusion carrying a sequential peptide affinity (SPA) tag at the C-terminus (Zeghouf *et al*., 2004). PipB2-SPA levels were reduced upon EsvB expression (Figure 5H). Northern blot analysis confirmed EsvB expression in the same samples (Figure 5H). Together, EsvB operates as a *cis*-encoded antisense sRNA, directly tuning SPI-2 effector levels thus affecting intracellular growth.

### EsvB targets genes encoding SPI-2 T3SS apparatus components associated with intracellular survival

Because macrophage survival assays showed that in trans ectopic expression of EsvB impaired the survival of wild-type *Salmonella* carrying an intact *pipB2* allele, as well as the Δ*pipB2* strain - indicating that *pipB2* is not the sole target of EsvB (Figure S10), we examined the SPI-2-encoded *ssaL*, another top predicted EsvB target and a component of the type III secretion system apparatus. Quantitative real-time PCR analysis revealed that P*lac*-EsvB repressed mRNA levels of *ssaH* and *ssaL*, the first and last genes of the operon, respectively, suggesting that EsvB affects expression of the entire operon encoding components of the T3SS apparatus (Figure 6A). To monitor the effect of EsvB on the expression of *ssaH* and *ssaL* in macrophages, cells were infected with wild-type *Salmonella* and the Δ*pipB2* strain, and chromosomal mRNA levels of *ssaL*, *ssaH*, and EsvB were quantified during infection (Figure 6BC). The results showed that, at later stages of infection, *ssaH* and *ssaL* mRNA levels were higher in the Δ*pipB2* strain compared to the wild type carrying an intact *pipB2* allele including EsvB, indicating EsvB modulates intercellular survival by downregulating *ssaH* and *ssaL* expression during late infection.

**Figure 6.**
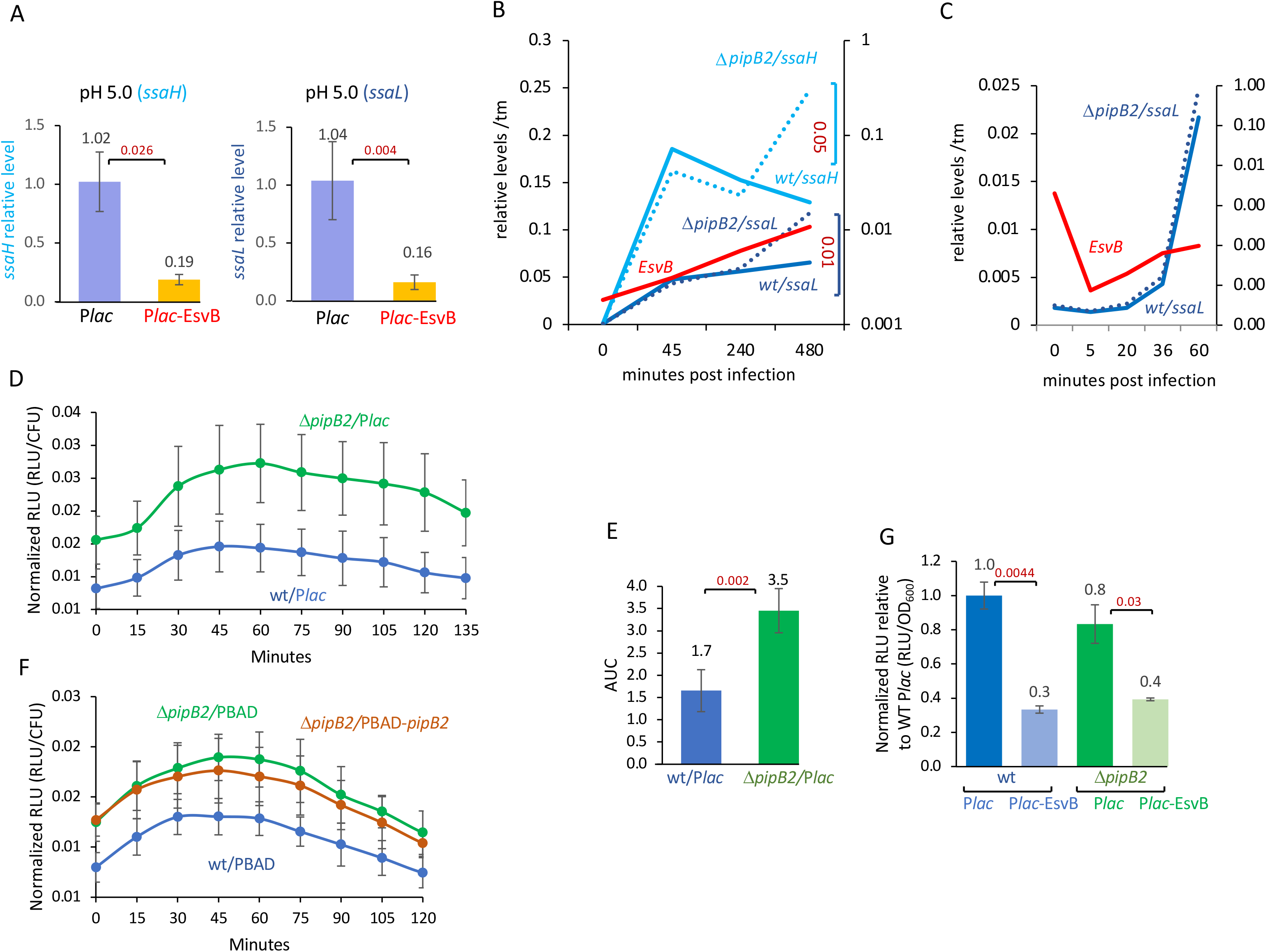
EsvB reduces secretion of SPI-2 T3SS (A) Quantitative real-time PCR analysis of *ssaH* and *ssaL* mRNA encoding proteins associated with the T3SS apparatus in the presence or absence of EsvB. Data represent mean ± SD of 3 biological replicates; *p*-values (two-tailed unpaired t-test) are indicated. (B, C) Macrophages were infected with *Salmonella* wild type (continuous line) and Δ*pipB2* (dashed line). At the indicated time points, samples were taken to measure chromosomal mRNA levels (RTPCR) of *ssaL*, *ssaH* and EsvB. *p*-values (two-tailed unpaired t-test) are indicated. (D) Kinetics of SopD2-HiBiT secretion in HeLa cells. HeLa LgBiT cells were infected with *Salmonella* wild type (blue) or Δ*pipB2* (green) as described in the methods. 5 hours post-infection, 0.2% arabinose to induce SopD2 expression and Nano-Glo substrate were added and luminescence was measured. In parallel CFU was calculated. Shown, kinetics of Relative Light Units (RLU) normalized to CFU of WT and Δ*pipB2* strains over time. (E) Area under the curve (AUC) calculated from normalized RLU data. Bars represent mean ± SD of 5 biological replicates. (F) Kinetics of SopD2-HiBiT secretion in HeLa cells following ectopic expression of PipB2. HeLa LgBiT cells were infected with *Salmonella* wild type (blue), Δ*pipB2* (green) or Δ*pipB2* with ectopic expression of *pipB2* deficient of EsvB. 5 hours post-infection, 0.2% arabinose was added to induce SopD2 expression and Nano-Glo substrate were added and luminescence was measured. In parallel CFU was calculated. Shown, kinetics of Relative Light Units (RLU) normalized to CFU over time. Bars represent mean ± SD of 3 biological replicates. (G) SopD2-HiBiT secretion into the medium was measured in wild-type and Δ*pipB2* strains carrying either a control plasmid or an EsvB-expressing plasmid. Cultures were grown for 4 hours in LB, then transferred to pH 5 medium for 6 hours in the presence of IPTG to induce EsvB expression. Samples were collected 30 minutes after the addition of arabinose to induce SopD2-HiBiT.

To assess the impact of EsvB-mediated repression of *ssaH-ssaL* operon on translocation of effectors, we analyzed SopD2 secretion. SopD2 is an SPI-2 T3SS-secreted effector that regulates the stability of *Salmonella* containing vacuoles (SCV), supports intracellular replication in macrophages, and contributes to virulence (Schroeder *et al*., D’Costa *et al*.). We measured translocation of a SopD2-HiBiT luminescent fusion protein into HeLa LgBiT cells infected with wild type and Δ*pipB2* strain deficient of both EsvB and PipB2 protein. Kinetics of SopD2-HiBiT secretion showed that SopD2-HiBiT secretion was higher in Δ*pipB2* strain than in wild type indicating that EsvB and/or PipB2 effector protein inhibited SopD2 secretion possibly by inhibiting expression of the operon encoding proteins responsible for T3SS apparatus (Figure 6DE). Secretion of SopD2-HiBiT following ectopic expression of PipB2-CDS deficient of EsvB sRNA (PBAD-*pipB2*) in Δ*pipB2* was comparable to that detected in Δ*pipB2* carrying the control plasmid, indicating that the secretion was not affected by the presence of PipB2 effector (Figure 6F). In addition, overexpression of EsvB inhibited SopD2-HiBiT secretion into the medium in either wild type or Δ*pipB2* (Figure 6G) further indicating that EsvB expression affects translocation mediated by SPI-2 Type III Secretion System apparatus.

### EsvB Couples SPI-1 Activation with SPI-2 Downregulation

EsvB targets genes and operons associated with the SPI-2 T3SS, thereby restricting intracellular survival. Intriguingly, CopraRNA predictions also identified *sipB*, an SPI-1 T3SS effector that promotes invasion rather than intracellular persistence as a potential target of EsvB (Table S3). This prompted us to examine the effect of EsvB on the *sicA*-*sipA* operon. Quantitative real-time PCR analysis of *sicA*, *sipB*, and *sipC*-representing the first, second and third genes of the operon-performed under high-density and pH 5 conditions showed that overexpression of EsvB increases mRNA levels of these genes under both conditions (Figure S11AB). The effect of EsvB on SicA protein levels was examined using a SicA-SPA fusion carrying a sequential peptide affinity (SPA) tag at the C-terminus (Zeghouf *et al*.). SicA-SPA levels increased ∼1.5-fold upon EsvB expression under pH 5 conditions, as quantified relative to total protein loading (Figure S11 CD). Together, these results indicate that EsvB exerts complementary effects: it promotes invasion while reducing survival within macrophages. This behavior is intriguing and suggests that EsvB coordinates opposing stages of *Salmonella* infection by balancing the transition between bacterial entry and intracellular survival.

## Discussion

### A high-throughput strategy for functional discovery of sRNAs

Landmark studies have cataloged bacterial transcripts (Kröger *et al*.), mapped RNA-RNA interaction networks (Melamed *et al*., Matera *et al*., Kooshapour *et al*.), and characterized infection-stage transcriptional programs (Nguyen *et al*.). However, a major bottleneck has remained, the systematic assignment of biological function to the large number of candidate regulatory RNAs uncovered by these approaches. By directly coupling an unbiased pooled sRNA expression library with a quantitative competition based functional readout, in this case an infection-based selection, our study provides a scalable solution to this challenge and reveals an unexpected layer of temporal regulation that coordinates distinct stages of *Salmonella* pathogenesis. This framework should be broadly applicable to functional discovery of regulatory RNAs in diverse bacterial systems.

The library captures a broad diversity of RNA species, including fragments originating from 5ʹ UTRs, coding regions, 3ʹ UTRs, antisense strands, intergenic regions, and known sRNAs. Interestingly, the largest fraction of fragments is derived from 5ʹ UTR regions, followed by CDS and 3ʹ UTRs, in-line with our current understanding of sRNA biogenesis, in which termination, internal promoters, and RNase-dependent processing generate abundant small RNAs from these genomic contexts (Ponath *et al*.). A substantial fraction (∼21%, ∼185 fragments) maps to antisense coding regions, as well as to the antisense strands of the 3ʹ and 5ʹ ends. Although many antisense fragments may reflect noise or artifacts (Thomason & Storz), ∼20 of these (Table S1) lie within 10 bp of transcription start sites or RNase E cleavage sites, suggesting authenticity.

Given that *Salmonella* virulence is governed by precise temporal regulation of its secretion systems across growth phases (Ilyas *et al*., Pérez-Morales *et al*., Bao *et al*., Li *et al*., Sia *et al*.) we compared the dataset of infection-permissive library to one generated from cells grown to high density (i.e., stationary phase). The two libraries showed similar distributions across genetic loci (Figure S12A); however, a notable difference was observed in the proportion of fragments originating from antisense loci (21% vs. 12%, respectively). It remains unclear whether this reflects reduced antisense transcription or decreased stability of the resulting fragments. Additionally, we observed differences in the enrichment of fragments derived from pathogenicity islands (Table S4). SPI-1, SPI-2, and SPI-5 were enriched in the infection-permissive library, whereas SPI-1, SPI-4, and SPI-5 were enriched in the high-density library. Given that SPI-1 drives invasion, SPI-2 supports intracellular survival, and SPI-4 mediates adhesion and cooperates with SPI-1 (Ilyas *et al*., Li *et al*.), the observed enrichment patterns i.e., SPI-2 under infection-permissive conditions and SPI-4 in the high-density library are unexpected. These results suggest that the library may capture early SPI-2 activation under infection-like conditions and SPI-4 persistence or delayed turnover during stationary phase. Alternatively, the patterns may reflect post-transcriptional regulation, including differences in stability or processing of SPI-derived fragments.

Conservation analysis showed that ∼50% of fragments in the infection-permissive library exhibit > 65% sequence conservation across Enterobacteriaceae, while ∼75% show at least 57% conservation among *Salmonella* subspecies. (Figure 1C and Figure S3). Conservation across evolutionary distances often suggests functional constraints, particularly for regulatory RNAs whose activity depends on conserved structural elements or base-pairing regions (Updegrove *et al*.).

RIL-seq, which captures the Hfq-mediated RNA-RNA interactome in *Salmonella* enterica (Matera *et al*.), was performed under high-density conditions and therefore shows greater overlap with the stationary-phase library (222 vs. 118 fragments) (Figure S12BD). Among fragments shared between RIL-seq and our datasets, most originated from 5ʹ regions (45% vs. 37% for infection-permissive and high-density conditions, respectively), followed by coding sequences (27% vs. 36.5%), sRNAs (12% vs. 9%), and 3ʹ ends (9% vs. 12.6%), supporting their authenticity. Approximately 24% of the fragments identified in macrophages overlapped with the Hfq-mediated RNA-RNA interactome (Matera *et al*.), indicating that a substantial fraction (∼25%) are functionally active in an Hfq-dependent manner (Figure S12CD).

The comparison between conventional RIL-seq performed under SPI2-inducing conditions (Kooshapour *et al*.) and our libraries (infection-permissive conditions, macrophage infection, and high-density library) indicates that 42%, 43%, and 48% of fragments are shared between our datasets and the RIL-seq data, respectively, indicating that our libraries may capture virulence-associated interactions comparable to those observed under SPI-2 inducing conditions. In contrast, only 14-18% of fragments are shared with intramacrophage RIL-seq dataset, which are less complex to begin with (Kooshapour *et al*.).

### Stage-specific sRNA regulation coordinates *Salmonella* infection dynamics

Functional screening further identified sRNAs with opposing effects on bacterial infection fitness, including two newly characterized 5ʹ UTR-derived regulators, EsvA and EsvB. Temporal profiling revealed a shift in sRNA activity, with some regulators limiting early stages of infection (e.g., EsvA), while others promote adaptation at later intracellular stages (e.g., Esv168, Esv232, and Esv252). Within this framework, based on their targets, EsvA appears to limit host cell entry, whereas EsvB promotes invasion while repressing SPI-2–dependent intracellular functions, suggesting complementary roles in coordinating transitions between infection stages. (see also model Figure 7).

**Figure 7.**
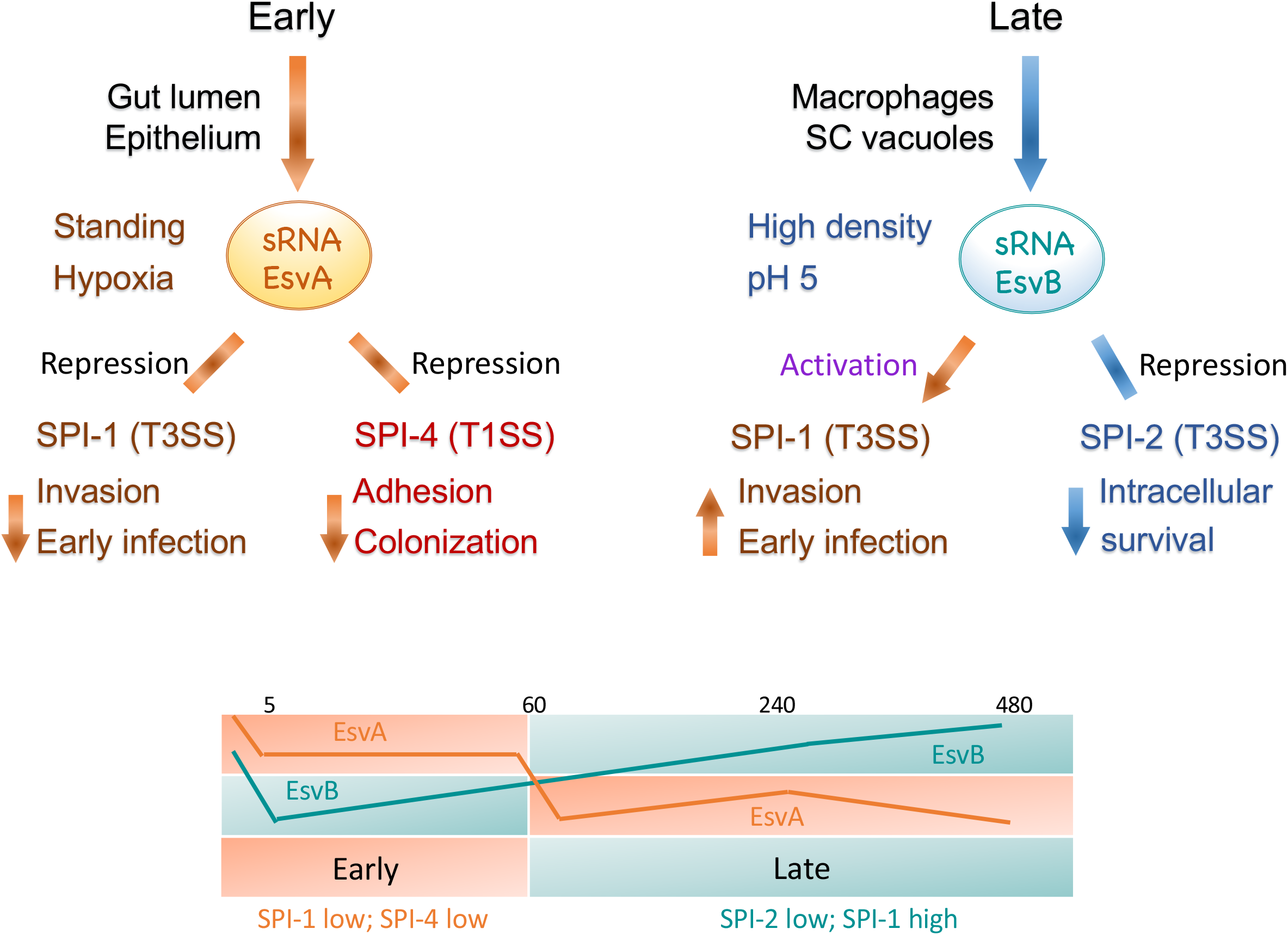
Stage-specific sRNA networks orchestrate *Salmonella* virulence (A) EsvA influences host-cell entry by regulating SPI-1 T3SS genes involved in invasion, as well as the SPI-4 operon associated with adhesion. In contrast, EsvB appears to coordinate the transition between host-cell entry and intracellular survival by activating SPI-1 while repressing SPI-2 T3SS expression. (B) EsvA predominates during the early stages of infection whereas EsvB becomes dominant at later stages indicating a regulatory switch between invasion and intracellular survival programs.

EsvA emerges as a multifaceted post-transcriptional regulator that modulates *Salmonella* virulence through coordinated regulation of multiple pathways associated with host cell entry. It stabilizes its parental transcript, STM0327, although the functional consequence of this effect remains unclear, while also repressing major virulence determinants within SPI-4 and SPI-1, including the *siiABCDEF* operon and the *orgABC* locus. These effects are associated with reduced adhesin production and altered secretion profiles. Together, these findings position EsvA as a modulator of early infection processes.

We propose that EsvA functions as a post-transcriptional stage-specific regulator that coordinates early stages of infection, during which precise control of secretion systems is critical for successful host colonization (Hautefort *et al*.). In this model, EsvA fine-tunes the transition between invasion and intracellular persistence by dampening early virulence gene expression (Figures 7 and S13).

In contrast, EsvB appears to orchestrate opposing virulence programs, acting as a regulatory switch between infection stages. It represses the expression of the SPI-2 effector *pipB2* and the *ssaH-ssaI* operon, potentially impairing intracellular secretion, while simultaneously activating SPI-1-associated genes. Despite enhanced SPI-1 expression, this does not translate into a detectable increased invasion, possibly owing to dominant suppression of SPI-2-dependent intracellular fitness or the limited sensitivity of the macrophage survival assay. Another possible explanation is that activation of SPI-1 in only a small bacterial subpopulation is sufficient to achieve maximal invasion (Sánchez-Romero & Casadesús). Notably, conventional RIL-seq performed under SPI-2-inducing conditions identified interactions involving *pipB2* (potentially mediated by EsvB) and the *ssaMVOT* operon, located immediately downstream of *ssaL–ssaH* (Kooshapour *et al*.), supporting a role for EsvB in modulating the SPI-2 apparatus. Together, these findings suggest that while EsvA modulates *Salmonella* virulence by coordinately regulating pathways involved in host-cell entry, EsvB may fine-tune the balance between invasion and intracellular adaptation, potentially preventing premature expression of intracellular functions.

Successful *Salmonella* infection requires the coordinated activation and repression of distinct virulence programs. Previous work established the canonical intergenic sRNA PinT as a post-transcriptional regulator that coordinates the transition between invasion and intracellular survival (Kim *et al*., Correia Santos *et al*., Kooshapour *et al*.). Whereas PinT exemplifies a canonical intergenic sRNA that post-transcriptionally coordinates virulence, EsvA and EsvB demonstrate that functionally important virulence regulators can also emerge from 5ʹ UTRs. Their discovery expands the repertoire of functional virulence regulators beyond canonical intergenic sRNAs and highlights non-canonical transcript-derived RNAs as an important and previously underappreciated layer of post-transcriptional control.

During the first hour of infection (Figure 7), EsvA expression remained relatively high following an initial decline, supporting a role in modulating early infection processes through the repression of SPI-1 and SPI-4 genes involved in invasion and adhesion (Main-Hester *et al*.). This ensures that host-cell entry is tightly controlled, preventing premature or excessive activation of invasion programs and reducing the fitness costs associated with virulence expression (Sturm *et al*.). As infection progresses, EsvA expression gradually decreases, relieving repression of these pathways and allowing the timely activation of virulence programs required for subsequent stages of infection. Collectively, this dynamic sRNA regulatory landscape enables *Salmonella* to optimize fitness by deploying specific virulence programs only when they are most advantageous for the current stage of the infection cycle. (Nguyen *et al*.).

### Limitations

It is important to note that construction of the library likely involves technical limitations inherent to high-throughput screening approaches, including biased cloning efficiencies, incomplete representation of all fragments, and the inability to clone unstable fragments. Therefore, we do not claim that the library provides a highly accurate quantitative “snapshot” of RNA coverage and/or abundance. Rather, we emphasize that the diversity and authenticity of the cloned fragments support the use of this library as a rich experimental platform for functional sRNA discovery, from which broad biological trends can be inferred. Several technical refinements could further improve library quality and accuracy. For example, more efficient rRNA depletion during library preparation would increase the effective diversity of regulatory RNA fragments. In addition, replacing the T1T2 transcriptional terminator with a self-cleaving ribozyme-based terminator (He *et al*.) could improve transcript boundary precision and reduce transcriptional read-through.

## Data availability

Sequencing data have been deposited in GEO under accession numbers GSE338675 and GSE338681

## Acknowledgments

The authors gratefully acknowledge Efrat Goloventzitz for her assistance with the analysis of the infection data, we thank Liron Argaman for her valuable advice on library construction, Prof. Retsef Levi for valuable discussions and constructive comments on the manuscript, Prof. Michael Hensel for providing the SiiE::HA strain and Prof. Ilan Rosenshine for providing us with the Hibit system and the MDCK cell line. We sincerely appreciate Rachel Marianovsky for her support in lab maintenance. This work was supported by the Israel Science Foundation founded by The Israel Academy of Sciences and Humanities (661/23). J.G. was supported by the DFG (GE 3159/1-1)

---

**Figure S1.**
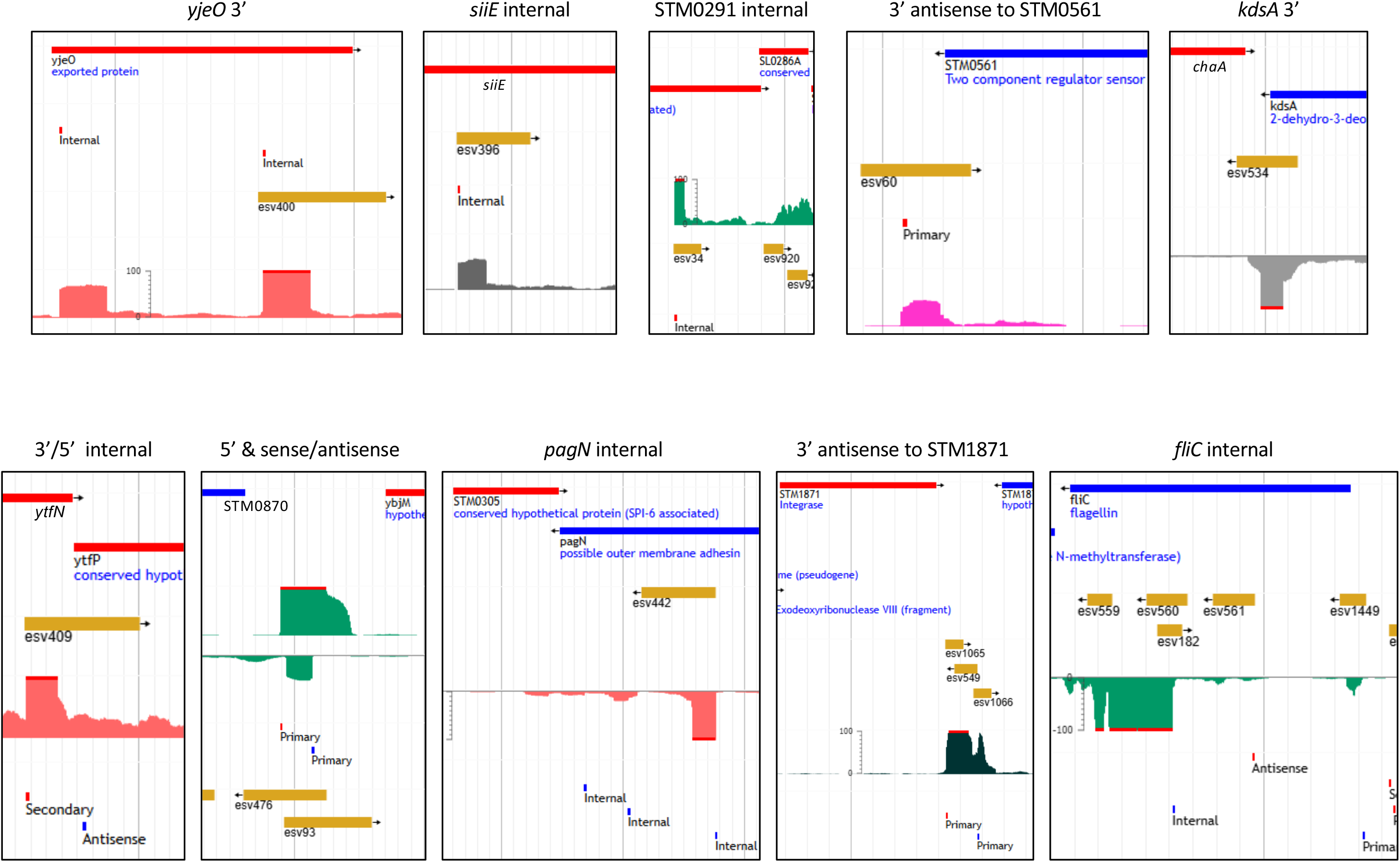
Library fragments (denoted esv for Experimentally discovered Small RNA associated with Virulence) were compared with existing Salmonella transcriptomic data [Kroger 2013]. Annotated genes, promoters (red: plus strand; blue: minus strand) and RNA expresion levels under varoius conditions are shown as reported in Kroger.

**Figure S2.**
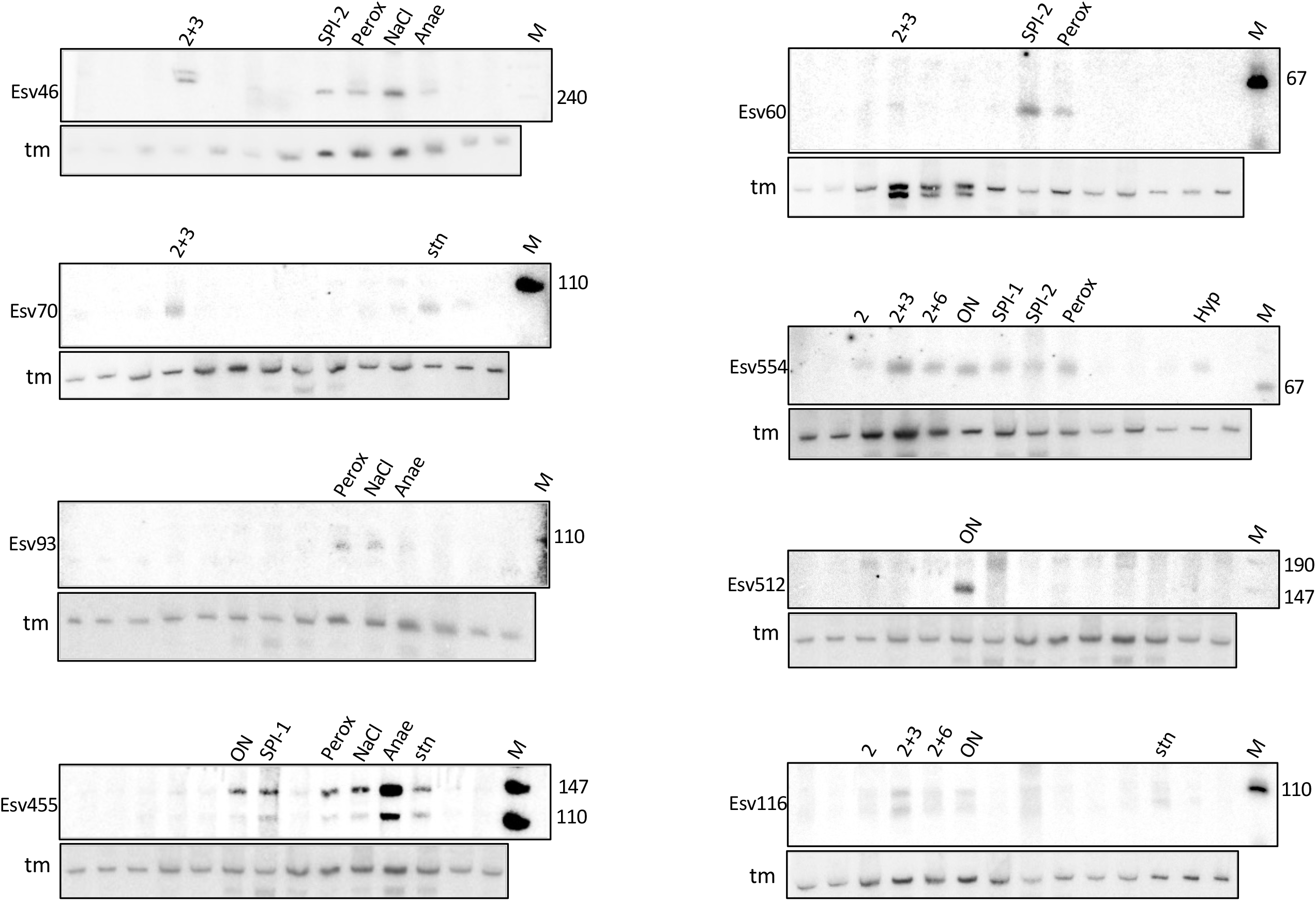
Northern blot validation of chromosomally encoded sRNA expression under defined conditions. tmRNA served as loading control (see also Figure 1B).

**Figure S3.**
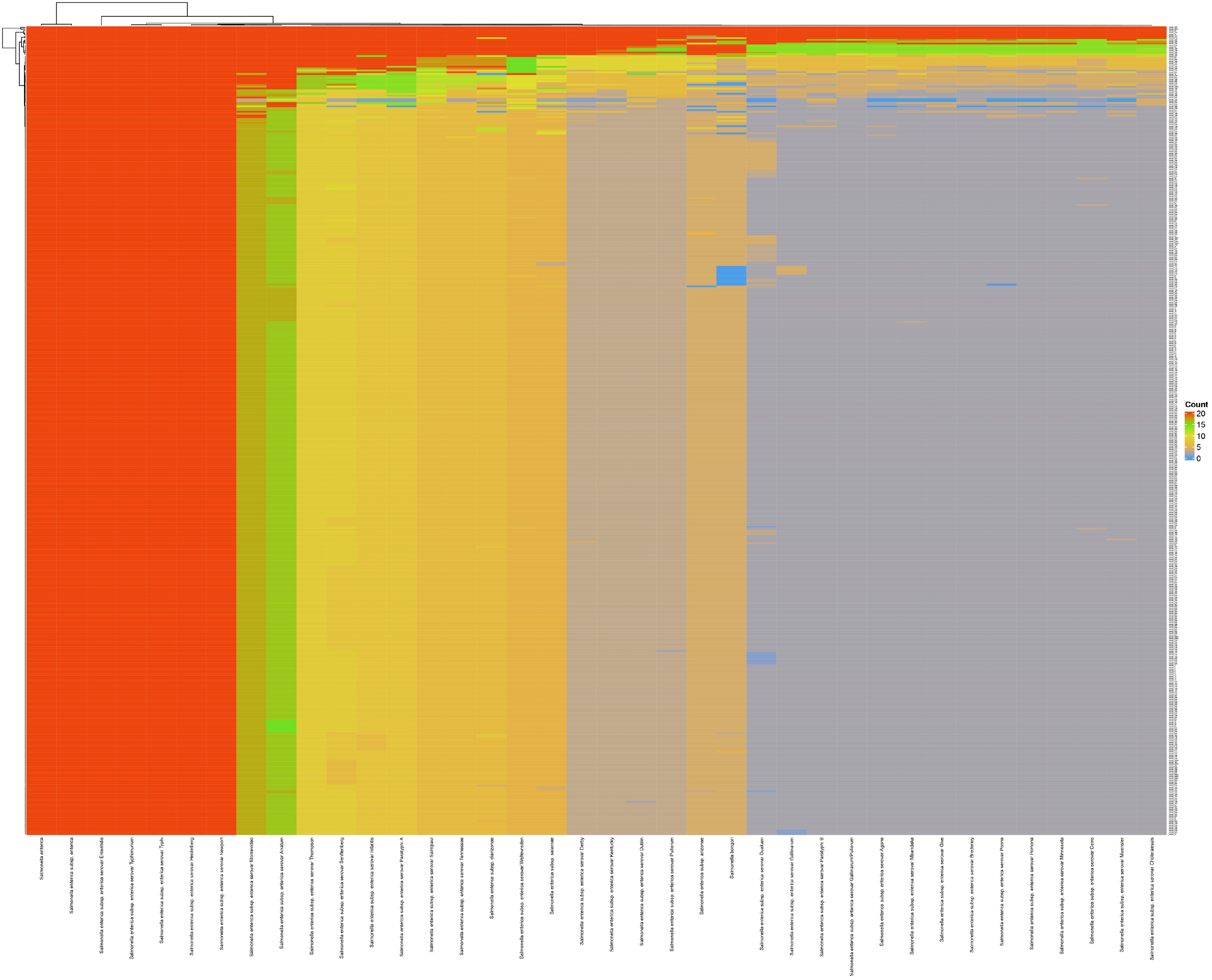
Conservation analysis of 450 fragments (minimum column sum ≥1000) are conserved (57%) within Salmonella subspecies, highlighting their potential functional relevance. Interactive Shiny-based maps were used for visualization. See also Figure 1C

**Figure S4.**
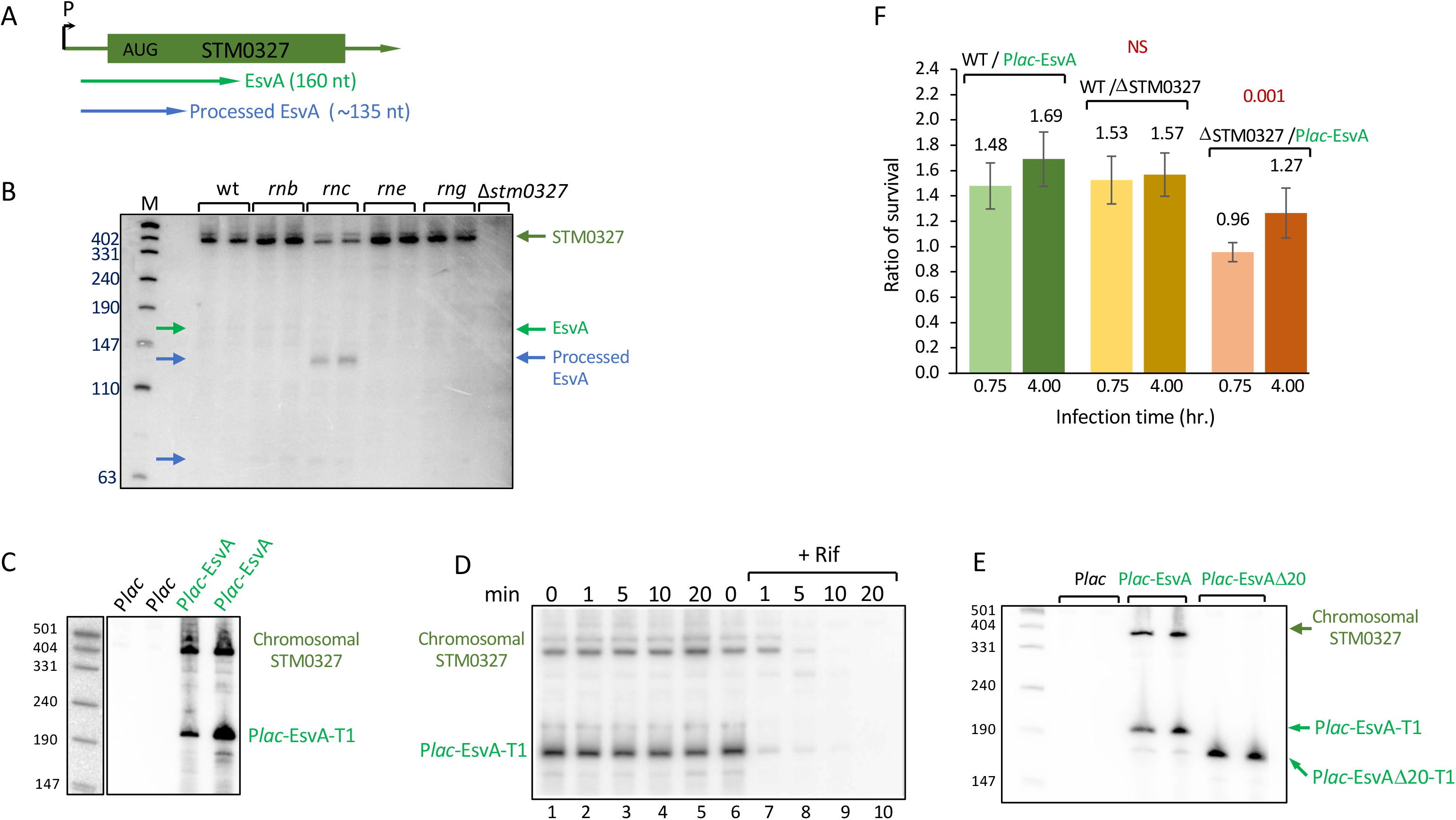
(A) Schematic representation of EsvA and STM0327, where EsvA originates from the 5ʹUTR of STM0327 and its position and orientation are indicated by the green arrow. (B) The stability of chromosomally encoded EsvA is positively influenced by RNase III (rnc) (C) Plasmid-encoded in trans expression of EsvA increases the transcript levels of the chromosomally encoded STM0327 (D) Rifampicin assay reveals that while EsvA RNA is highly unstable (lanes 6 and 7), it stabilizes STM0327 mRNA. In the absence of EsvA, the level of STM0327 is drastically reduced. (E) An EsvA mutant with an internal 20-nucleotide deletion (see Figure 3D) failed to stabilize STM0327 mRNA. Panels B, C, D and E display northerns data. (F) Time-course survival comparisons: wild-type relative to EsvA overexpression (green), wild-type relative to the STM0327 mutant (brown), and STM0327 mutant relative to EsvA overexpression (orange).

**Figure S5.**
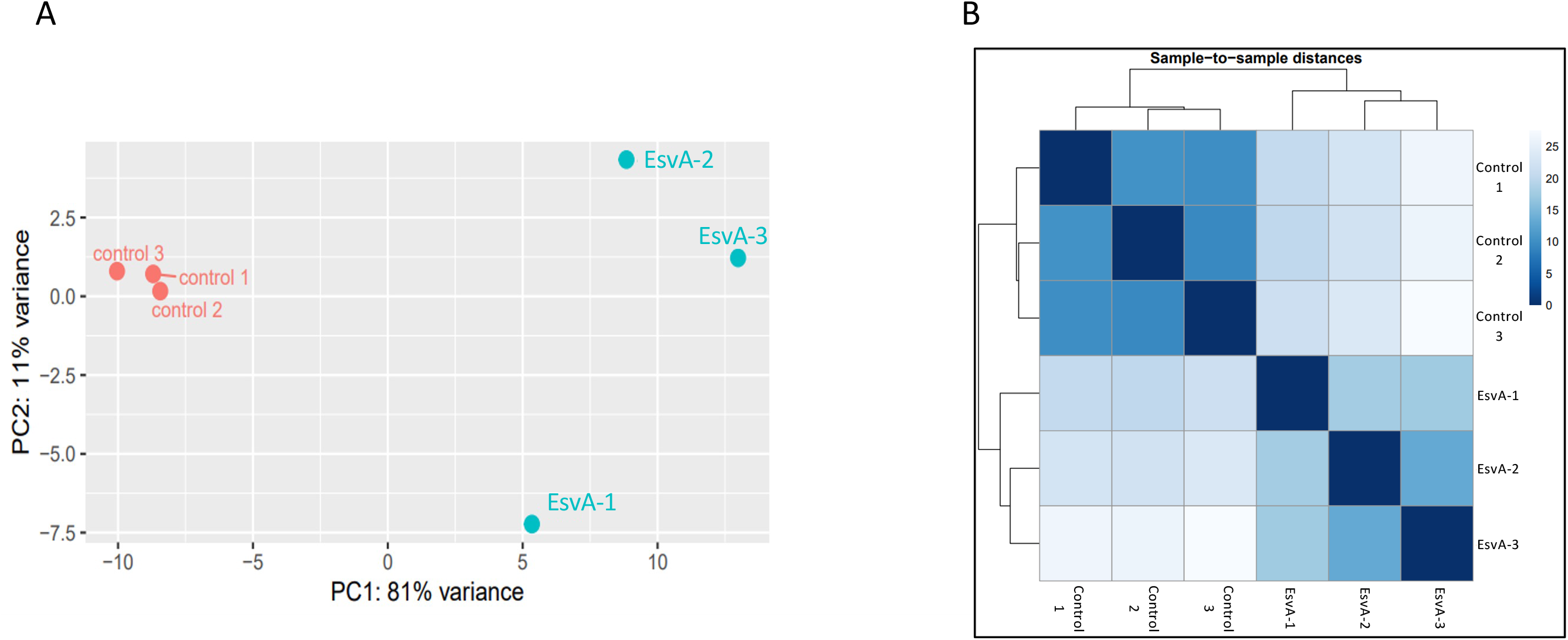
(A) Principal component analysis (PCA) at 30 min comparing EsvA overexpression and control samples, with separation primarily along PC1 (81% variance explained) and PC2 (11% variance explained (B) Distance heatmap at 30 min showing clustering of three biological replicates of EsvA overexpression and three replicates of control samples.

**Figure S6.**
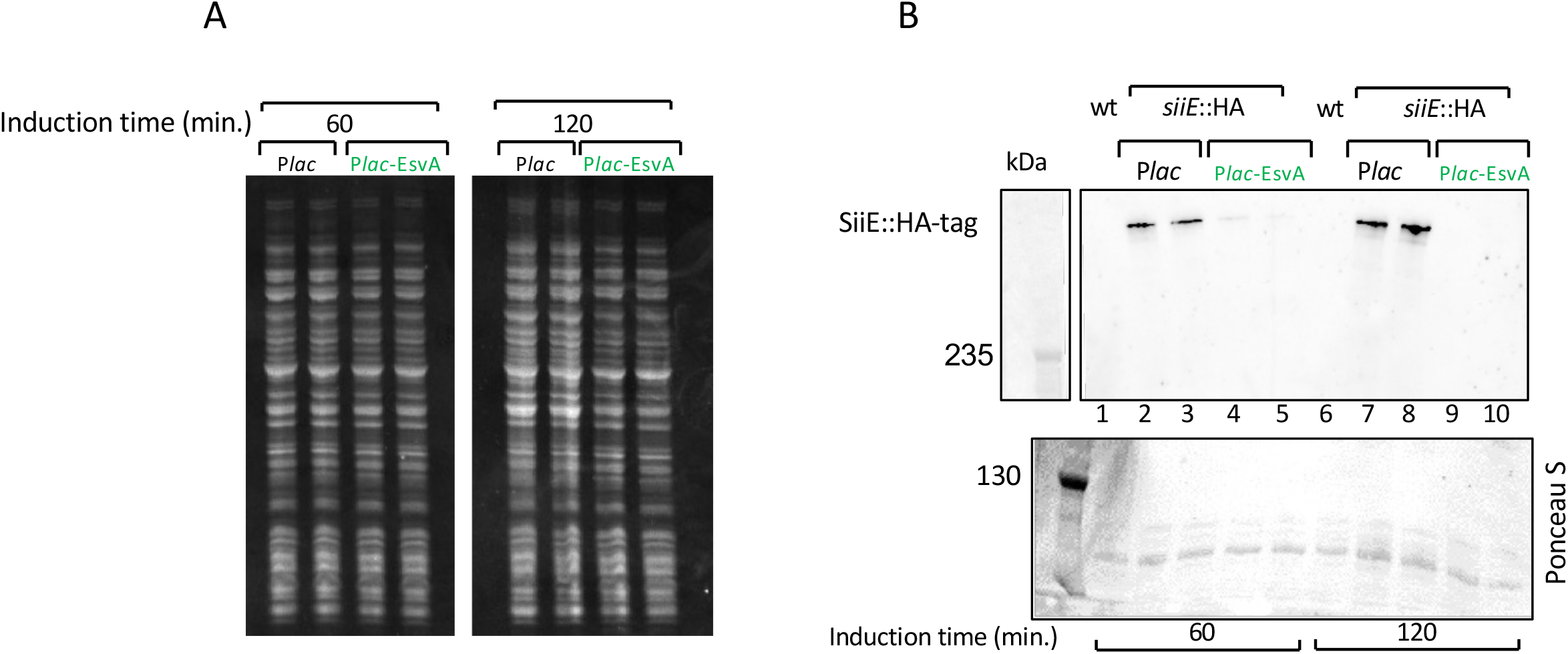
(A) Coomassie blue staining of samples collected at 60 and 120 min after IPTG induction for subsequent western blotting. (B) A strain carrying tagged SiiE::HA shows reduced protein levels upon *in trans* expression of EsvA. Western blot (upper panel). Ponceau S staining (lower panel) was used as loading control. Wild type (wt) carries the native, non-engineered allele of *siiE* and serves as a control.

**Figure S7.**
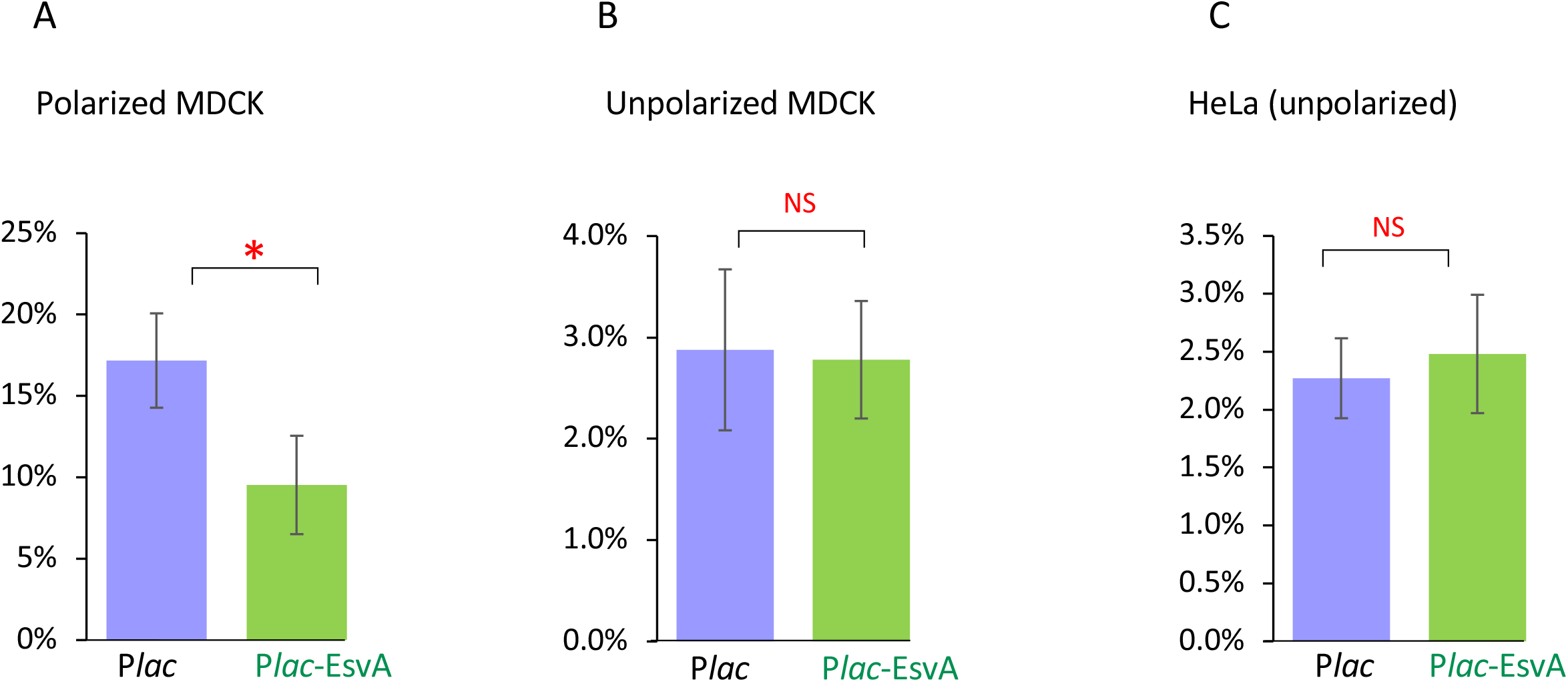
Adhesion assays show that *in trans* ectopic expression of EsvA impairs the adhesion of *Salmonella* to polarized MDCK cells and not to unpolarized cells. (A) Adhesion percentage to 6 days polarized MDCK cells. (B-C) Adhesion percentage to unpolarized MDCK cells and HeLa cells. Data represent mean ± SD of 4-5 biological replicates; p-values (two-tailed unpaired t-test) are indicated.

**Figure S8.**
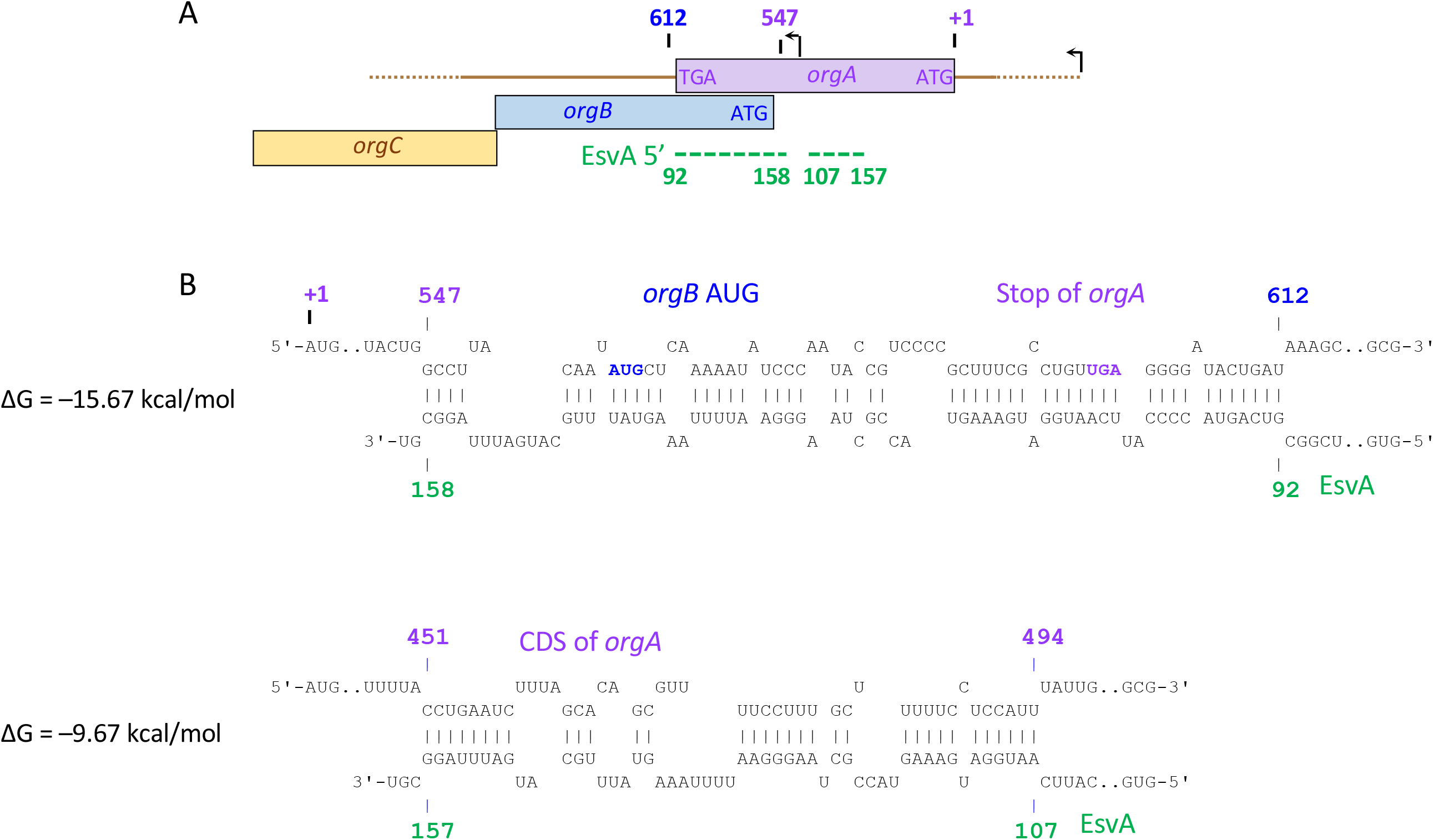
Predicted RNA - RNA interaction between the *orgABC* locus and EsvA, as calculated using the IntaRNA algorithm. Two EsvA molecules are predicted to bind distinct regions within *orgAB*. Position +1 corresponds to the AUG start codon of *orgA*. Arrows indicate promoters: one that directs transcription of the entire *prgHIJK–orgABC* operon, and another that specifically drives transcription of *orgBC*.

**Figure S9.**
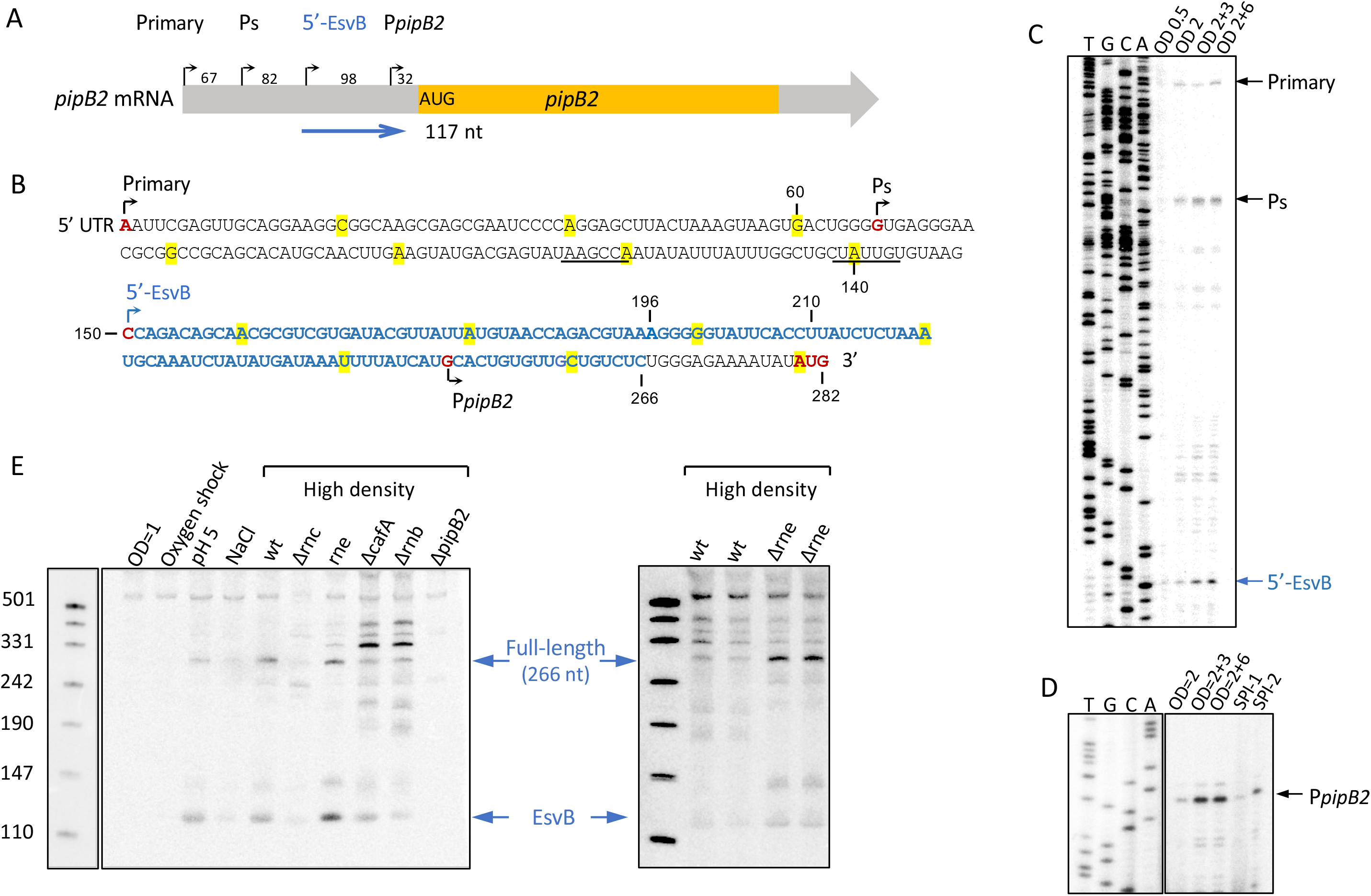
(A) Schematic representation of the pipB2 transcript showing three previously identified promoters: the primary promoter (Primary), a secondary promoter (Ps), and one located immediately upstream of pipB2 (PpipB2). The blue arrow indicates the position and orientation of EsvB. (B) Nucleotide sequence of EsvB (blue). The putative −10 and −35 elements of 5ʹ-EsvB are underlined, and the initiation codon of pipB2 is highlighted in red. (C–D) Transcriptional mapping of pipB2. Primer extension was performed using 20 μg of total RNA and end-labeled pipB2-specific primers 3957 (C) and 4116 (D). (E) Northern blot detecting short and full-length EsvB transcripts, the latter generated from the primary promoter. “High density” refers to cultures harvested at OD 2+3 h. The *rne* mutant strains used in this experiment are listed in the list of strains.

**Figure S10.**
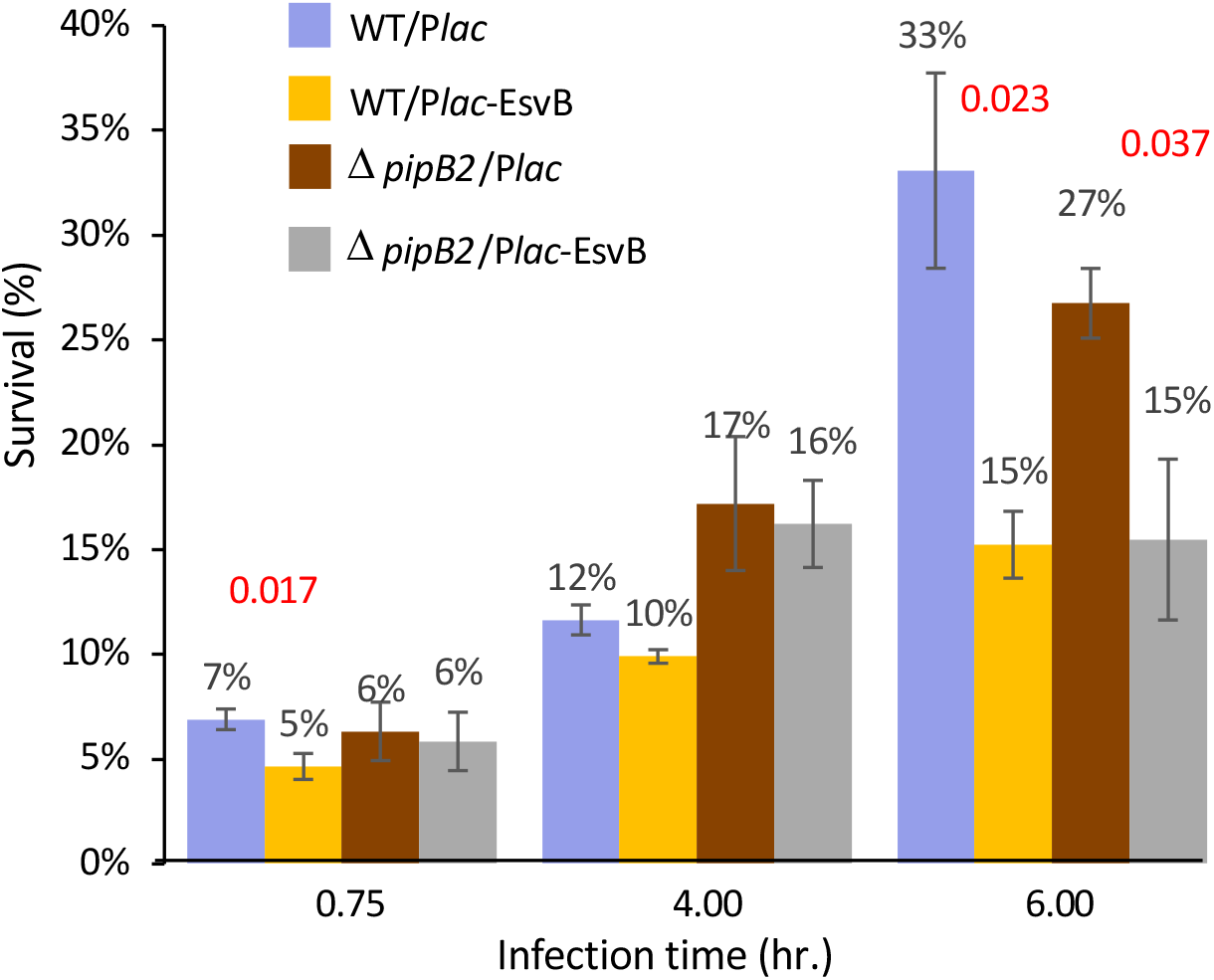
Macrophage survival assays show that *in trans* ectopic expression of EsvB impairs the survival of wild-type *Salmonella* carrying an intact *pipB2* allele, as well as that of the Δ*pipB2* strain. This indicates that *pipB2* is not the only target of EsvB. Data represent mean ± SD of 3-6 biological replicates; p-values (two-tailed unpaired t-test) are indicated.

**Figure S11.**
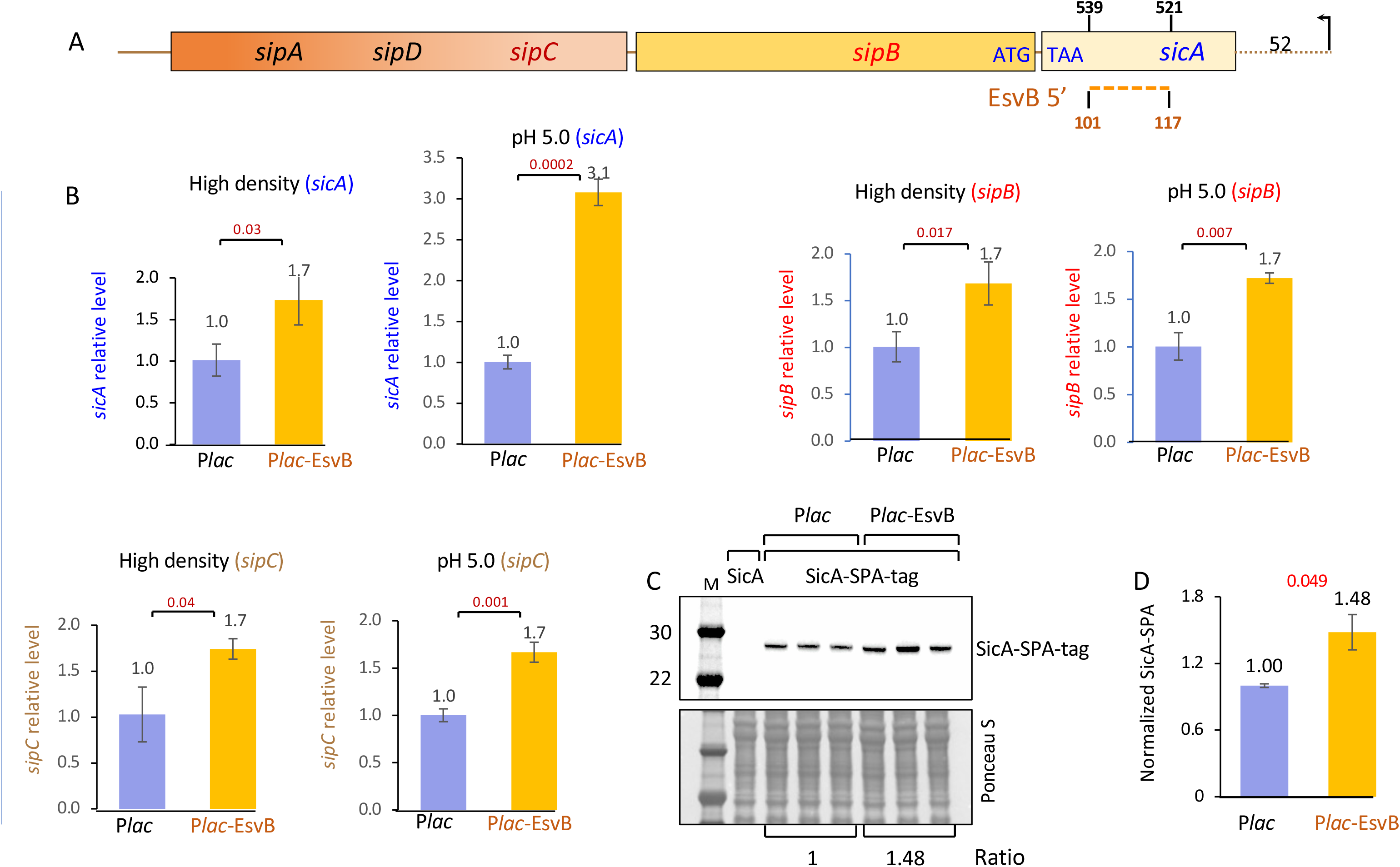
*In-trans* ectopic expression of EsvB reduces *pipB2* mRNA levels and increases *sicA sipB sipc* mRNA levels under both high cell density conditions and at pH 5.0. (A) Schematic representation of the *sicA, sipB, sipC, sipD, sipA* locus, encompassing SPI-1 type III secretion-associated proteins involved in bacterial entry. Possible RNA-RNA interaction between the *sicA* and EsvB, as predicted using the IntaRNA algorithm (red dashed line). (B Quantitative real-time PCR analysis of *sicA sipB sipC* expression in the presence or absence of EsvB. Data represent mean ± SD of 3 biological replicates; p-values (two-tailed unpaired t-test) are indicated. (C) Western blot of SicA-SPA expression in the presence of EsvB and its quantification (D) Cultures of wild type and SicA-SPA were grown for 4 hours in LB, then transferred to pH 5 medium for 6 hours in the presence of IPTG to induce EsvB expression. Wild type lacking SPA carrying P*lac* served as control.

**Figure S12.**
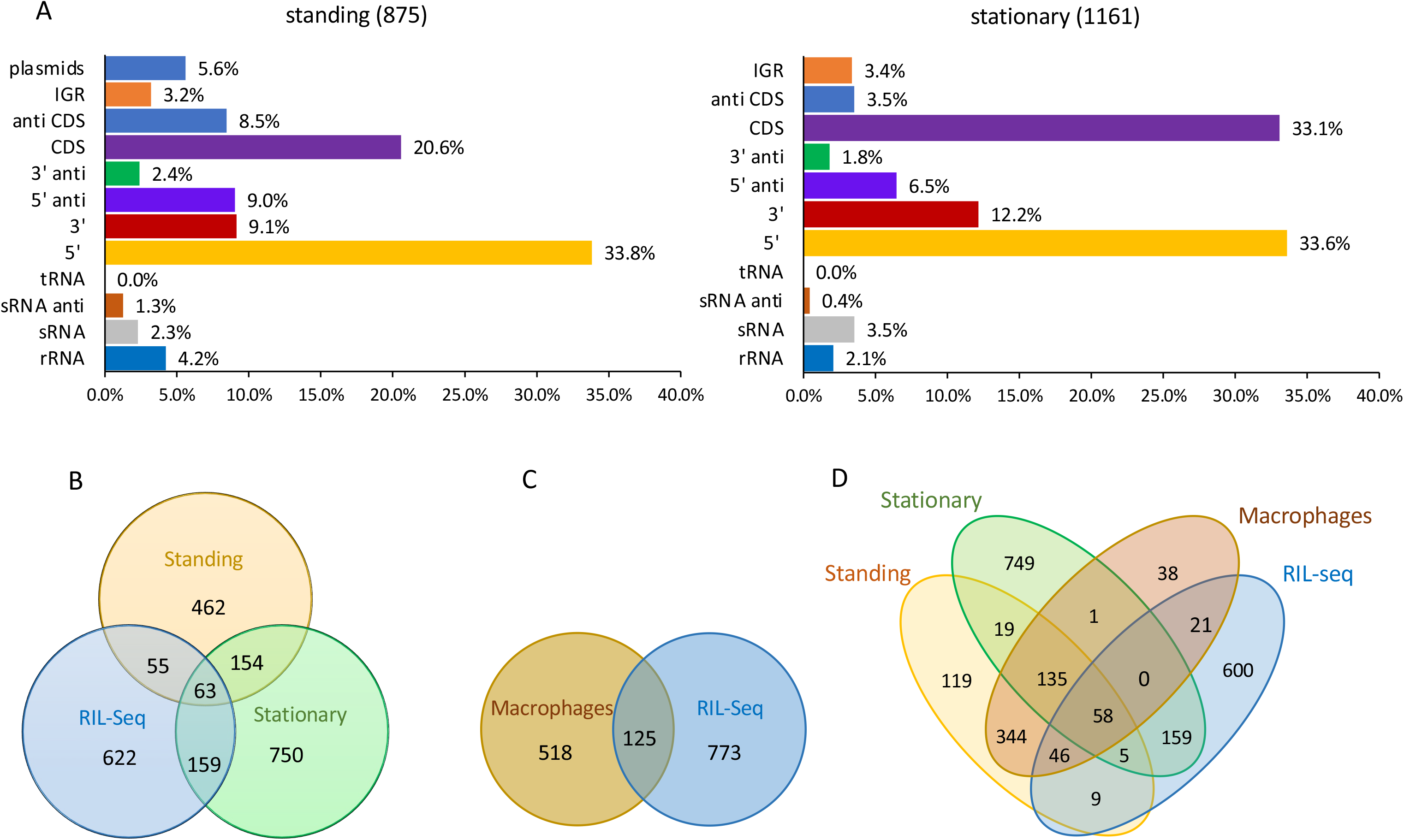
Fragments shared between libraries collected under standing and stationary conditions and in RIL-seq data. (A) Genomic distribution of fragments collected under standing and stationary conditions. (B) Overlapping fragments between the two libraries and the published RIL-seq data (C) Overlapping fragments collected upon macrophage infection and RIL-Seq data. (D) Overlapping fragments collected upon macrophage infection RIL-Seq data and the two libraries. Published RIL-seq data was generated from cultures grown to OD600 of 2.0. RIL-seq fragments were calculated by adding up to 200 bp from the start of the first RNA read. rRNA and tRNA were excluded from these analyses.

**Figure S13.**
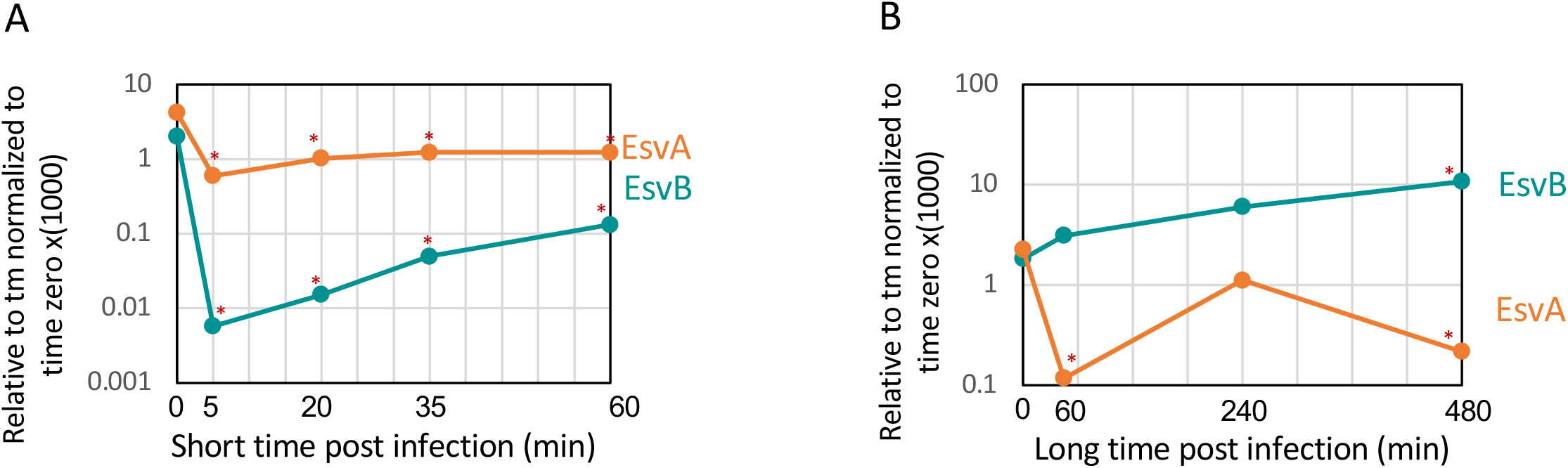
EsvA predominates during the early stages of infection, (A) whereas EsvB becomes dominant at later stages (B), indicating a regulatory switch between invasion and intracellular survival programs. Macrophages were infected with wild-type *Salmonella*. At the indicated time points, samples were collected to measure chromosomal EsvA and EsvB transcript levels by RT-PCR. Values were calculated relative to tmRNA, normalized to time zero, and multiplied by 1000. P values (two-tailed unpaired *t* test) are indicated. Asterisks denote significance relative to time zero.

**Table S4:** sRNA enrichment in genomic islands: stationary vs. standing.

| Island | Size | Stationary library |  |  |  | Standing library |  |  |  |
| --- | --- | --- | --- | --- | --- | --- | --- | --- | --- |
|  |  | No. of sRNAs<sup>b</sup> | Ratio<sup>c</sup> | Library ratio<sup>d</sup> | Enrichment<sup>e</sup> | No. of sRNAs<sup>b</sup> | Ratio<sup>c</sup> | Library ratio<sup>d</sup> | Enrichment<sup>e</sup> |
| SPI-1 | 42204 | 48 | 1E-03 | 2.2E-04 | 5.1 | 15 | 4E-04 | 1.5E-04 | 2.4 |
| SPI-2 | 40165 | 5 | 1E-04 | 2.2E-04 | 0.6 | 10 | 2E-04 | 1.5E-04 | 1.7 |
| SPI-3 | 17022 | 2 | 1E-04 | 2.2E-04 | 0.5 | 2 | 1E-04 | 1.5E-04 | 0.8 |
| SPI-4 | 24750 | 22 | 9E-04 | 2.2E-04 | 4.0 | 4 | 2E-04 | 1.5E-04 | 1.1 |
| SPI-5 | 9109 | 7 | 8E-04 | 2.2E-04 | 3.4 | 7 | 8E-04 | 1.5E-04 | 5.1 |
| SPI-6 | 46314 | 9 | 2E-04 | 2.2E-04 | 0.9 | 7 | 2E-04 | 1.5E-04 | 1.0 |
| SPI-9 | 16681 | 0 | 0E+00 | 2.2E-04 | 0.0 | 0 | 0E+00 | 1.5E-04 | 0.0 |
| SPI-11 | 8413 | 3 | 4E-04 | 2.2E-04 | 1.6 | 2 | 2E-04 | 1.5E-04 | 1.6 |
| SPI-12 | 5266 | 0 | 0E+00 | 2.2E-04 | 0.0 | 0 | 0E+00 | 1.5E-04 | 0.0 |
| partial SPI-13 | 7478 | 1 | 1E-04 | 2.2E-04 | 0.6 | 0 | 0E+00 | 1.5E-04 | 0.0 |
| SPI-14 | 7372 | 3 | 4E-04 | 2.2E-04 | 1.8 | 2 | 3E-04 | 1.5E-04 | 1.8 |
| SPI-16 | 4198 | 0 | 0E+00 | 2.2E-04 | 0.0 | 0 | 0E+00 | 1.5E-04 | 0.0 |
| SLP105 | 45242 | 5 | 1E-04 | 2.2E-04 | 0.5 | 3 | 7E-05 | 1.5E-04 | 0.4 |
| bacteriophage SLP203 | 40088 | 11 | 3E-04 | 2.2E-04 | 1.2 | 5 | 1E-04 | 1.5E-04 | 0.8 |
| Oaf O-antigen modification locus | 9905 | 0 | 0E+00 | 2.2E-04 | 0.0 | 1 | 1E-04 | 1.5E-04 | 0.7 |
| CS54 genomic island | 24139 | 2 | 8E-05 | 2.2E-04 | 0.4 | 2 | 8E-05 | 1.5E-04 | 0.6 |
| SLP272 | 50513 | 13 | 3E-04 | 2.2E-04 | 1.2 | 11 | 2E-04 | 1.5E-04 | 1.5 |
| degenerate bacteriophage SLP281 | 10534 | 2 | 2E-04 | 2.2E-04 | 0.9 | 0 | 0E+00 | 1.5E-04 | 0.0 |
| SLP285 | 32907 | 7 | 2E-04 | 2.2E-04 | 1.0 | 6 | 2E-04 | 1.5E-04 | 1.2 |
| prophage SLP289 | 11907 | 6 | 5E-04 | 2.2E-04 | 2.3 | 3 | 3E-04 | 1.5E-04 | 1.7 |
| prophage remnant SLP443 | 22114 | 2 | 9E-05 | 2.2E-04 | 0.4 | 2 | 9E-05 | 1.5E-04 | 0.6 |

<sup>a</sup>Size of the island in bases<sup>b</sup>Number of short RNA fragments identified within an island<sup>c</sup>Ratio of short RNA fragments to island size<sup>d</sup>Ratio d Ratio of total short RNA fragments to total genome- 1088/4878013 and 732/4878013 for stationary and standing libraries, respectively.<sup>e</sup>Enrichment - ratio of c to d (rRNA and tRNA were excluded. Areas with enrichment higher than 1.5 are highlighted in red).

## Materials and Methods

### Bacterial growth conditions

*Escherichia coli* and *Salmonella* cultures were grown at 37°C (200 rpm) in LB medium (pH 6.8). Ampicillin (100 μg/ml), tetracycline (10 μg/ml), chloramphenicol (20 μg/ml), and kanamycin (40 μg/ml) were added where appropriate. P*lacO* promoter was induced with isopropyl β-D-thiogalactoside (IPTG) 0.4 mM unless otherwise specified, PBAD promoter was induced with 0.2% arabinose unless otherwise specified. For RNA extraction and sRNA identification, *Salmonella* lacI^q^ strain were grown under various physiological conditions. In most cases, cultures were grown overnight in 3 ml LB medium at 37°C with shaking at 200 rpm, followed by a 1:100 dilution into fresh medium. Cultures were grown until reaching an OD_600_ of 0.15, 0.3, 0.5, 2.0, and late stationary phases (3 and 6 hours after OD_600_ = 2.0) (Kröger *et al*.). For overnight condition, cultures were grown for 18 hours after dilution. For SPI-1 inducing conditions, single colonies were inoculated into 5 ml LB supplemented with 0.3 M NaCl in closed 15 ml tubes and grown for 12 hours at 37°C with shaking. For SPI-2 inducing conditions, cultures were grown overnight in PCN medium (pH 5.8, 0.4 mM Pi) and diluted 1:100 into fresh PCN medium until reaching an OD_600_ = 0.3 (Kröger *et al*.). For anaerobic shock, cultures grown to OD_600_ = 0.3 in LB were transferred to tight 50 ml tubes and incubated without agitation for 30 minutes (Kröger *et al*.). Hypoxia shock was induced similarly by shifting OD_600_ = 0.45 cultures to closed tubes for 4 hours (Avican *et al*.). Oxidative stress (peroxide shock) was induced by adding 1 mM H_2_O_2_ to OD_600_ = 0.3 cultures in PCN medium for 12 minutes (Kröger *et al*.). Osmotic shock was induced by adding 0.3 M NaCl to OD_600_ = 0.3 cultures grown in LB for 10 minutes (Kröger *et al*.). For intra-cellular conditions, RAW264.7 macrophages were infected at a MOI of 1:50 using bacterial cultures grown 18 hours at 37°C without shaking (standing condition). Infection was carried out as described at macrophage growth conditions and macrophages survival assay sections. Samples were collected 45 minutes, 4 hours, and 8 hours post-infection.

### Macrophage growth conditions

RAW 264.7 macrophages were regularly grown at 37°C and 5% CO_2_ in DMEM high glucose media (Sigma, cat. no. D5796) containing 10% heat-inactivated fetal-bovine-serum (Biological Industries, cat. no. 04-127-1A) supplemented with Penicillin-Streptomycin (Biowest, cat. no. L0022). 24 hours prior to infection, the medium was replaced by Penicillin-Streptomycin free medium. Appropriate antibiotics, IPTG and arabinose were added to the macrophage’s cultures before infection. Macrophages were activated by adding 6μg/ml phorbol 12-myristate-13-acetate (PMA) right before infection with bacteria cells.

### HeLa growth conditions

HeLa cells were regularly grown at 37°C and 5% CO_2_ in DMEM high glucose media (Sigma, cat. no. D5796) containing 10% heat-inactivated fetal-bovine-serum (Biological Industries, cat. no. 04-127-1A) supplemented with Penicillin-Streptomycin (Biowest, cat. no. L0022). On the same day of infection, the medium was replaced by Penicillin-Streptomycin free medium. Appropriate antibiotics, IPTG and arabinose were added to the cultures before infection.

### MDCK growth conditions

MDCK cells were regularly grown at 37°C and 5% CO_2_ in DMEM high glucose media (Sigma, cat. no. D5796) containing 10% heat-inactivated fetal-bovine-serum (Biological Industries, cat. no. 04-127-1A) supplemented with Penicillin-Streptomycin (Biowest, cat. no. L0022). 4 to 10 days before infection, 1 x 10^6^ cells were seeded in a 24 wells plate. 4 hours prior to infection medium was replaced by Penicillin-Streptomycin free medium. Appropriate antibiotics and IPTG was added to the cultures before infection. For polarized MDCK, cells were grown for 6 days.

### Strain construction

Chromosomal gene deletion mutants were carried out using the chloramphenicol cassette of pKD3 (Datsenko & Wanner, Yu *et al*.) on SL1344 ΔhisG::(PlacI^q^-lacI-Tn10, Tet^r^) strain. To construct Δ*pipB2*::Cm, primers 4148 and 4149 were used to replace *pipB2* genomic region, from P1 promoter, with chloramphenicol cassette. To construct ΔSTM0327::Cm, primers 4227 and 4228 were used to replace ∼0.5 kb region encompassing STM0327 with chloramphenicol cassette. To generate Δ*pipB2*::FRT the antibiotic resistance genes were removed using pCP20 (Datsenko & Wanner). Chromosomal SPA tagging were carried out using the SPA-kanamycin cassette of pJL148 (Zeghouf *et al*.) on SL1344 ΔhisG::(PlacI^q^-lacI-Tn10, Tet^r^) strain. To construct *pipB2*-SPA-*kn*, primers 4110 and 4111 were used. To construct *orgB-SPA-kn*, primers 4356 and 4357 were used. To construct *sicA-SPA*-*kn*, primers 4383 and 4389 were used.

### Plasmid construction

To isolate specific plasmids from the library, primers located in the center of the insert were used, and the plasmid containing the specific insert was amplified using whole plasmid PCR. After phosphorylation of the PCR product, the plasmid was re-circularized with T4 DNA ligase (New England BioLabs, cat. no. M0202S). The primers used for each construct were as follows: for P*lacO*-Esv252-t1t2, 3769 and 3770; for P*lacO*-EsvB-t1t2, 3956 and 3957; for P*lacO*-EsvA-t1t2, 3958 and 3959; for P*lacO*-Esv168-t1t2, 4171 and 4172; and for P*lacO*-Esv232-t1t2, 4169 and 4170. To construct PBAD-STM0327, primers 4094 and 4120, were used to amplify the gene from SL1344 genome. The PCR product was digested with the PstI and HindIII restriction enzymes and ligated into similarly digested pEF21 (Guzman *et al*.). To construct *pipB2*-*lacZ* translation fusion, 356bp upstream of *pipB2* AUG and 32pb downstream were PCR amplified using primers 4120 and 4187. The PCR products were digested with KpnI and BamHI and ligated into similarly digested pBOG552 plasmid (Hershko-Shalev *et al*.). To construct *pipB2-lacZ* 4M (DS site) and *pipB2-lacZ* 3M (US site) mutants, the KpnI and BamHI fragment was inserted into pGEM3 plasmid and subjected to whole plasmid PCR using primers 4190–4191 and 4229-4230, respectively. After ligation, the mutated fragment was then amplified from the pGEM3 plasmid using primers 4126 and 4187, digested with KpnI and BamHI and ligated into similarly digested pBOG552 plasmid. To construct P*lacO-siiA*-*lacZ* translation fusions, sequences were PCR amplified using primers #4279-#4280. The PCR products were digested with KpnI and BamHI and ligated into similarly digested pBOG552-P*lacO* plasmid. To construct *sopD2-FLAG-HiBit* translation fusion, 54bp upstream of *sopD2* AUG until last codon were PCR amplified using primers 4367 and 4368. The PCR products were cloned, using Gibson assembly (Gibson *et al*.), into pSA10-HiBit plasmid linearized with whole plasmid PCR with primers 4369 and 4370. *sopD2-FLAG-HiBit* fragment was PCR amplified using primers 4385 and 4386. The PCR products were digested with PstI-HF and ligated into similarly digested pEF21 plasmid (Guzman *et al*.). Cloned plasmids were checked for correct insert orientation using primers 1314 and 4372 and Sanger sequencing. To construct pBAD-*pipB2*, 26bp upstream of *pipB2* AUG and 119pb downstream to the stop-codon were PCR amplified using primers 4109 and 4409. The PCR products were digested with EcoRI and HindIII and ligated into similarly digested pJO244 plasmid (Hershko-Shalev *et al*.).

### Library construction

#### RNA extraction and size selection

Total RNA was isolated from SL1344 WT strain grown overnight under infection permissive conditions (‘standing’) or to stationary phase (OD_600_ = 2) using TRI-reagent (Sigma, cat. no. T9424). Following extraction, the RNA (100 µg) was treated with KAPA pure beads (Cat no. KK8000) (0.65X to 3.5X volumetric ratio) followed by Zymo RCC-25 (Cat. no. R1017) to keep RNA fragments ranging from 100 nt to 450 nt. The resuspended RNA (in 5 µl) was loaded onto acrylamide gel (6%, 8M urea) and fragments ranging from 50 nt to 400 nt were excised and eluted from the gel by adding 2 volumes/weight of elution solution (500 mM NH4OAc, 1 mM EDTA-KOH pH8) followed by overnight incubation at 4°C. The eluted RNA was extracted using phenol-chloroform followed by ethanol precipitation. To remove DNA, 2.2 µg RNA was treated with 10 U of Turbo DNase (Thermo Fisher Scientific, cat. no AM2238) 40 U RRI (Takara, cat. no. 2313 A) and 1X DNase buffer and incubated at 37°C for 30 min followed by Zymo OCC (Cat. no. D4060).

#### Adapters ligation, cDNA synthesis and cloning

RNA (11-15 pmol) was ligated with adenylated 3’ adapter (3233) (20-30 pmol) in a reaction mixture carrying 300 U of truncated KQ T4 RNA ligase (New England BioLabs, cat. no. M0373), 15% PEG8000, 12 U of RRI and 1X T4 RNA ligase buffer (4 hours at 16°C). Ligase was inactivated using 5 mM EDTA, and the reaction was purified with KAPA pure beads (3.5X volumetric ratio) and eluted with 12µl. For cDNA synthesis, 100 pmol of primer 3239 (complementary to 3’ adapter) and 100 pmol of primer 3397 (complementary to pBR plac0-1 promoter) were added to the ligated RNA and the reaction was incubated for 3 min at 72°C and then for 2 min at 42°C. Thereafter, the resulting RNA was subjected to reverse transcription reaction (in 200 U of SMARTscribe (Clontech, cat. no. 639538), 2.5X SMARTscribe buffer, 6.25 mM DDT, 2.5 mM dNTP and 20 U RRI) for 90 min at 42°C, followed by inactivation at 70°C for 10 min. The cDNA product was purified using KAPA pure beads (3X volumetric ratio), followed by 9 or 15 cycles of amplification using KAPA HIFI HotStart ready mix (Cat. no. KK2602) and 3398 and 3239 primers. The PCR product was purified with KAPA pure beads (2.5X volumetric ratio) and cloned into pBR-PlacO-t1t2 using Gibson assembly (4 hours at 50°C) (Gibson *et al*.) with 1:4 to 1:10 ratio of PCR product to plasmid. The product was purified using Wizard SV Gel and PCR Clean-Up System (Promega, cat. no. A9281), introduced into XL1-Blue cells and the transformants were plated on 35 plates (LB containing ampicillin). The resulting colonies were scraped from plates with 10 ml fresh LB. 10 samples of 400µl each were subjected for plasmid extraction using a QIAprep Spin Miniprep Kit (Cat. no. 27104). The rest of the resuspended colonies were saved at -80°C.

#### Library sequencing

Cloned fragments were subjected to 15 cycles of amplification from the plasmid library, using backbone primers with Illumina overhangs adapters (3581 and 3582) and KAPA HIFI HotStart ready mix. The PCR product was purified and primers were removed by using KAPA pure beads (3X volumetric ratio), and concentration was measured using Qubit. Second PCR, using Illumina DNA Prep, and Next-generation sequencing, using NextSeq 500/550 Mid-Output v2.5 Kit (150 cycles) (Cat. no. 20024904), was carried out by the Genomic Applications lab, Hebrew University of Jerusalem.

#### library analysis

For the initial cluster definition fastQ files (GSE338675) from the stationary and infection-permissive libraries were concatenated, the reads were quality trimmed, filtered and adapters were cut with fastp (https://doi.org/10.1002/imt2.107). Remaining reads were mapped to the Salmonella enterica subsp. enterica serovar Typhimurium SL1344 and its plasmids (FQ312003.1, HE654724.1, HE654725.1, HE654726.1) with segemehl (https://doi.org/10.1186/gb-2014-15-2-r34). Individual cloned RNA segments were identified using BlockClust (10.1093/bioinformatics/btu270). For the initial cluster definition fastQ files (GSE338675) from the macrophage infection timeseries were concatenated, the reads were quality trimmed, filtered and adapters were cut with fastp (https://doi.org/10.1002/imt2.107). Remaining reads were mapped to the Salmonella enterica subsp. enterica serovar Typhimurium SL1344 and its plasmids (FQ312003.1, HE654724.1, HE654725.1, HE654726.1) with segemehl (https://doi.org/10.1186/gb-2014-15-2-r34). Individual cloned RNA segments were identified using BlockClust (10.1093/bioinformatics/btu270). The resulting 717 clusters from the macrophage dataset were merged with 875 and 1161 clusters from a standing condition and a stationary phase condition library, respectively to yield 1754 non-overlapping clusters. The cluster-based annotation was used to count the reads in the individual macrophage infection timeseries datasets with htseq-count (10.1093/bioinformatics/btu638) prior to differential expression analysis with DESeq2 (https://doi.org/10.1186/s13059-014-0550-8).

#### Annotation of expressed cloned RNA clusters

For a systematic annotation the genomic positions of the RNA clusters were compared with the annotated features on the Salmonella enterica str. SL1344 genome (FQ312003.1) and the plasmids pSLTSL1344 (HE654724.1), pCol1B9SL1344 (HE654725.1) and pRSF1010SL1344 (HE654726.1). A match within a genomic feature was assigned to this respective feature, e.g. CDS, ncRNA, rRNA or tRNA and 5’ UTRs defined based on manually curated dRNAseq data (Kröger *et al*.) using the closest available TSS to each gene. A match within 100nt downstream of an annotated CDS was assigned as 3’UTR. The same logic applied to the assignment of clusters antisense to UTRs or genomic features. Additionally, the clusters were matched with Salmonella pathogenicity islands SPIs (Kröger *et al*.) and novel small protein CDSs (Venturini *et al*.). Next the start coordinates of the cloned RNA clusters were compared with known transcriptional start sites (TSS) (Kröger *et al*.) and RNase E cleavage sites (Chao *et al*.). A file with all annotations is available as Supplementary Table S1. The clusters were assigned to the feature using the following hierarchy: sRNA > 5’UTR > 3’UTR > CDS > rRNA > tRNA > anti-SRNA > anti-5’UTR > anti-3’UTR > anti-CDS > IGR.

#### Phylogenetic distribution expressed cloned RNA clusters

A GLASSgo (https://doi.org/10.3389/fgene.2018.00124) homolog each was done for each clustered library RNA fragment against the NCBI nucleotide (nt) blast database (release dated: Jan 26, 2018) for minimum percentage identity of 52 with structural clustering (londen mode) turned off. Taxonomic information for all included organisms was extracted from the NCBI taxonomy resource (https://www.sciencedirect.com/science/article/pii/S1673852721000837) with TaxonKit. A shiny app (https://github.com/JensGeorg/sRNA_distribution_heatmap) allows the visualization of the distribution of all or selected clusters within selectable organism lists and/or taxonomic groups. A web-based version of the app can be accessed via shinyapps.io (https://jensrna.shinyapps.io/sRNA_conservation_heatmap/). Comparative sRNA target prediction for esvA and esvB CopraRNA prediciton for esvB was done using 11 and for esvA using 16 sRNA homologs respectively. The full results and involved organisms are available as Supplementary Table S2 (EsvA) and S3 (EsvB).

#### Northern blot analysis

RNA samples (7-30 µg) isolated from strains as indicated were denatured for 10 min at 70°C in 98% formamide loading buffer, separated on 6% acrylamide 8 M urea gels and transferred to BrightStar®-Plus membranes (Applied Biosystems) by electroblotting. To detect the sRNA and target genes, the membrane was hybridized with end-labeled primers (table X) in modified CHURCH buffer (1 mM EDTA, pH 8.0, 0.5 M NaHPO_4_, pH 7.2, and 5% SDS) for 2 h at 45°C and washed as previously described (Ben-Zvi *et al*.). Tm RNA (10Sa) was used as a loading control.

#### Macrophages selection screen

The library of plasmids was transformed, in three duplicates, into SL1344 lacI^q^, and after 1 hour of recovery, the transformed cells were moved into 5 ml of fresh LB media containing selective antibiotic (Ampicillin) for overnight growth at 37°C with shaking. The cultures were then diluted 1:100 in 30 ml of fresh LB media containing selective antibiotic and grown for 18h at 37°C without shaking (‘standing’). sRNAs expression was induced with 1mM IPTG after 18h for 1h. Based on OD 600, an appropriate volume to reach 1:50 MOI was used to infect activated, macrophages grown as described above in 150mm dishes (Thermo Scientific, cat. no. 168381). After 30 minutes of incubation at 37°C and 5% CO_2_ incubator, dishes were washed twice with pre-warmed 1X PBS, and fresh media containing 100µg/ml gentamicin was added to kill all extracellular bacteria. Cultures were then incubated for 45 minutes, washed twice with warm 1X PBS, and from 10 dishes (for every duplicate) plasmid was extracted by adding 1X PBS supplemented with 1% triton X-100 followed by scraping and extraction with QIAprep Spin Miniprep Kit (Cat. no. 27104). To the rest of the dishes fresh medium containing 100µg/ml gentamicin was added, the cultures were incubated for an additional 7 hours upon which 10% triton X-100 was added directly to dishes (final 1%) and plasmids were extracted as previously.

Extracted plasmid were subjected to 15 cycles of amplification, using backbone primers with Illumina overhangs adapters (3581 and 3582) and KAPA HIFI HotStart ready mix (Cat. no. KK2602). The PCR products were purified and primers were removed using AMPure XP beads (1.8X volumetric ratio) (Beckman Coulter, cat. No. A63881). The quality and size of PCR products were assessed using Tapestation (Agilent Technologies, high sensitivity D1000 kit). Second PCR, using Illumina DNA Prep, and Next-generation sequencing, using NextSeq 500/550 High-Output v2.5 Kit (75 cycles) (Cat. no. 20024906), was carried out by the Genomic Applications lab, Hebrew University of Jerusalem.

#### Macrophages survival assays

A day prior to infection RAW 264.7 macrophages were harvested from a T75 flask by gentle scraping, transferred to 50 ml tubes and centrifuged at room temperature for 3 minutes at 1000 rpm. The cells pellet was resuspended in appropriate volume of fresh DMEM Penicillin-Streptomycin free medium, and homogenized by pipetting. Cells concentration was determined and the final concentration was adjusted to seed 0.8-1.1 × 10⁶ cells per well in 1 ml of medium in a 24-wells culture plates (Thermo Scientific, cat. no. 142475). Cells were incubated overnight at 37°C in a humidified 5% CO₂ incubator. Right before infection with bacteria cells 6 μg/ml PMA, 1 mM IPTG were added to the macrophage cultures. Bacteria cultures of SL1344 lacI^q^, SL1344 lacI^q^ Δ*pipB2*::FRT and SL1344 lacI^q^ ΔSTM0327::FRT carrying P*lac* plasmids, as indicated, were prepared by diluting 1:100 of overnight cultures in 10 ml of fresh LB media containing appropriate antibiotics. The cultures were grown at 37°C with shaking (200 rpm) for 4h (OD_600_∼ 2) upon which, 400 µL were transferred into Eppendorf tubes and resuspended in 800 µL of fresh LB medium. OD_600_ was measured and the required volume of bacterial suspension was used to infect the macrophages (MOI 1:10) and for plating on LB plates (to recalculate accurate CFU used for infection). Infected cultures were incubated at 37°C, 5% CO₂ for 30 minutes to facilitate infection. Following incubation, wells were washed twice with 1 ml warm 1X PBS, and fresh medium (DMEM, 10% FBS, 1 mM IPTG, 100 µg/mL gentamicin) was added to eliminate extracellular bacteria and induce sRNA expression. After 45 minutes, wells were harvested for bacterial quantification, and the remaining wells were washed twice again with 1 ml warm 1X PBS, followed by the addition of fresh medium containing low concentration of gentamicin (DMEM, 10% FBS, 1 mM IPTG, 10µg/mL gentamicin). Plates were incubated at 37°C, 5% CO₂ and harvested at indicated time points post-infection. Harvesting was done by washing the wells twice with 1 ml of 1X PBS, followed by adding 0.5 ml of PBS containing 1% Triton X-100. The lysate was pipetted up and down several times to ensure complete cell lysis, transferred to glass tubes and vortexed for 10 seconds, followed by incubation at room temperature for 10 minutes. Lysate was serially diluted with 1X PBS and appropriate amounts were plated on LB plates. CFU was counted and survival was calculated.

#### Total RNA-sequencing (EsvA, STM0327)

Three duplicates of SL1344 lacI^q^ carrying (1) P*lac* and PBAD (p15A) (2) P*lac-*EsvA and PBAD (p15A) (3) P*lac* and PBAD*-*STM0327 were grown from 1:100 dilution to OD_600_ of 0.2 in 125 ml flask. Thereafter, the cultures were exposed to 1 mM IPTG and 0.2% arabinose for 30 min at which total RNA was extracted with TRI-reagent (Sigma, cat. no. T9424). 400 ng RNA (Nanodrop) were subjected to fragmentation in FastAP buffer at 92°C for 1.5 min. DNA digestion and dephosphorylation were carried out simultaneously by adding 10 U of FastAP (Thermo Fisher Scientific, cat. no. EF0651), 8U of Turbo DNase (Thermo Fisher Scientific, cat. no. AM2238) and 40U of RRI (Takara, cat. no. 2313 A), at 37°C for 30 min. RNA samples were purified with Zymo RCC-5 (Cat. no. R1015), eluted with 12µl of which 5µl were incubated for 2 min at 70°C with 100pmol of 5’ phosphorylated (barcoded) adapters (primers 83, 84, 116, 117, 118, 119, 120, 121, 122). Thereafter, 14 µl of ligation mix (51,000 U of T4 RNA ligase 1 (New England Biolabs, cat. No. M0437M), 12.8% DMSO, 1.42X ligase buffer, 1.4 mM ATP, 28.5% PEG 8000 and 12 U of RRI) were added and the reactions and were incubated at 22°C for 2 hours. The reactions were stopped by the addition of 60 µl of RLT buffer (Qiagen, cat. no. 79216). The barcoded samples were pooled, purified with Zymo RCC-5 and eluted with 15 µl of DEPC. rRNA was depleted using Ribo-Zero gram-negative kit (Illumina, cat. no. MRZGN126, standard protocol). The samples were purified with 2.5X volumetric ratio of AMPure XP beads (Beckman Coulter, cat. No. A63881) and eluted with 12 µl of which 11 µl were used to generate cDNA by adding 50 pmol of AR2 primer (82). The mixture was incubated for 2 min at 70°C prior to the addition of reverse-transcription mix containing 200 U of SMARTscribe RT enzyme (Clontech, cat. No. 639538), 2.5 mM dNTP, 2.5X SMARTscribe buffer, 5 mM DTT and 20 U of RRI. The reaction was incubated further at 42°C for 1 hour, followed by inactivation for 15 min at 70°C. RNA was degraded by adding 2.5 µl of 1N of NaOH and incubation for 12 min at 70°C. Fresh acetic-acid was added (final concentration of 90 mM) and DEPC was added to the final vol of 40 µl. Primers were removed with 2.5X volumetric ratio of AMPure XP beads and eluted with 6 µl without removing the beads. 80 pmol of 3Tr3 adapter (85) was added to the RNA-beads mix, incubated for 3 min at 75°C and a ligation mix was added, containing 45,000U of T4 RNA ligase 1, 6.1% DMSO, 1.53X ligase buffer, 1.5 mM ATP and 32.6% PEG 8000 and the reaction was incubated overnight at 22°C. Reaction volume was adjusted to 40 µl, purified twice with 2.5X volumetric ratio of AMPure XP beads and eluted with 25 µl. cDNA product was subjected to 12 cycles of amplification using KAPA HIFI HotStart ready mix (Cat. no. KK2602) and primers containing Illumina P5 and P7 overhangs (86 and 87, respectively). The PCR products were purified with AMPure XP beads (1.5X volumetric ratio) and eluted with 12 µl. Sequencing of the samples were carried out at the Genomic Applications lab, Hebrew University of Jerusalem using NovaSeq 6000 (50M reads, Single-end, 50 cycles, cat. no. 20028314)

#### RNA-sequencing analysis (EsvA, STM0327)

Reads were quality trimmed (>25), demultiplexed and adapters were cut with Cutadapt (https://doi.org/10.14806/ej.17.1.200). 12nt form the head were trimmed and only reads with minimal length of 15nt were kept using Trimmomatic (https://doi.org/10.1093/bioinformatics/btu170). Remaining reads were mapped to the *Salmonella* enterica subsp. enterica serovar Typhimurium str. ST4/74 genome (NC_016857.1) and SL1344 plasmids (NC_017720.1, NC_017718.1, NC_017719.1) with Bowtie2 (https://doi.org/10.1038/nmeth.1923). Features were counted using featureCounts (https://doi.org/10.1093/bioinformatics/btt656) prior to differential expression analysis with DESeq2 (https://doi.org/10.1186/s13059-014-0550-8). To identify genes uniquely differentially expressed by EsvA, genes altered by STM0327 over-expression were subtracted from the differentially expressed genes in the EsvA over-expression dataset. Data is available at GSE338675.

#### Quantitative Real-Time PCR

For RNA isolation from cultures grown in LB media, overnight cultures of SL1344 lacI^q^ carrying P*lac* plasmids, as indicated, were diluted 1:100 and grown to the indicated growth condition in LB media: For over-expression of EsvA cultures were grown to OD_600_ = 0.4 and 0.4 mM of IPTG was added 30 minutes prior to harvesting. For high-density over-expression of EsvB cultures were grown to OD_600_ = 2 and then for additional 3 hours, IPTG was added after dilution. For pH 5 (Gong *et al*.) over-expression of EsvB cultures were grown for 4 hours, centrifuged at room-temperature and resuspended with fresh LB pH 5 media (adjusted with HCl) supplemented with 0.4 mM of IPTG, 6 hours prior to harvesting.

For intracellular RNA isolation, OD_600_ = 2 bacterial cultures were used to infect RAW 264.7 macrophages grown in 6-wells culture plates, after activation with PMA at a MOI of 1:10. For short-term kinetics, 100 µg/ml gentamicin was added to cell wells after 15 minutes and intracellular bacteria were harvested at 5, 15, 30, and 60 minutes post-infection. For long-term kinetics, after 30 minutes of infection, wells were washed twice with 1X PBS and fresh media containing 100 µg/ml gentamicin was added. After 45 minutes wells were washed twice again with 1X PBS and fresh media containing 10 µg/ml gentamicin was added. Intracellular bacteria were harvested at 0.75, 4 and 8 hours post-infection. At each indicated time point, wells were washed twice with warm 1X PBS, and macrophages were lysed using 1% Triton X-100 in PBS. The resulting lysate was vortexed vigorously for 10 seconds, and a fractional aliquot was removed for CFU quantification. The remaining bacterial fraction was pelleted by centrifugation at 4°C, resuspended in 50 µl TE buffer supplemented with 5 µl Lysozyme, and subjected to three rapid liquid nitrogen freeze-thaw cycles to ensure efficient bacterial cell wall disruption.

Total RNA was extracted using TRI-reagent (Sigma, cat. no. T9424). To maximize recovery from intracellular extraction, 1.5µl of GlycoBlue 15 mg/mL (Invitrogen cat. no. AM9515) was added during the isopropanol precipitation step, followed by an overnight incubation at -20°C. To ensure high purity, the RNA underwent a secondary ethanol precipitation (0.1 vol sodium acetate, and 2.5 vol ethanol) overnight at -20°C prior to concentration measurement. 2µg of clean RNA was subjected to DNase treatment using RQ1 DNase (Promega, cat. no. M6101, 2 units). cDNA synthesis was carried out using MMLV-RT (Promega, cat. no. M170A) and hexamer random primers (Promega, cat. no. C1181). Relative quantification was carried out using the Pfaffl Method (Pfaffl). For *siiF*, *siiA*, *orgA, orgB, orgC, pipB2, ssaH, ssaL, EsvA, EsvB, sicA, sipB* and *sipC*, tmRNA was used as a reference gene. For *<u>EsvB</u>* accurate quantification, the number of EsvB-*pipB2* reads was subtracted from *EsvB* reads.

#### In vitro RNA synthesis

DNA templates for RNA synthesis: *pipB2* (200 nt) was generated using primers 4111 and 4050; *siiA* (262 nt) was generated using 4298 and 4280; EsvA sRNA (160 nt) was generated using primers 4300 and 4301. The RNAs were synthesized in 50 µL reactions containing T7 RNA polymerase (25 units; New England Biolabs, cat. no. M0251), 40 mM Tris–HCl (pH 7.9), 6 mM MgCl_2_, 10 mM dithiothreitol (DTT), 20 units RNase inhibitor (TAKARA, cat. no. # 2313A), 500 µM of each NTP, and 200 ng of purified PCR templates carrying the sequence of the T7 RNA polymerase promoter. Synthesis was allowed to proceed for 2 h at 37°C followed by 10 min at 70°C. To remove the DNA template, 4 U of turbo DNase I (Ambion) was added (37°C, 30 min), followed by phenol/chloroform extraction and ethanol precipitation in the presence of 0.3 M ammonium acetate.

### In vitro RNA-RNA binding

#### siiA

Annealing mixtures containing: 0.05 pmol of in vitro-synthesized *siiA* RNA, without or with increasing concentrations of in vitro-synthesized EsvA RNA and 0.6 pmol of end-labeled *siiA*-specific primer (4280) were incubated for 10 min at 70°C followed by 10 min on ice. The mixtures were incubated for another 1 min at RT in 20 mM Tris–HCl, 10 mM magnesium acetate, 0.1 M NH_4_Cl, 0.5 mM EDTA, 2.5 mM β-mercaptoethanol (Sigma, cat. no. 8057400250), and 0.5 mM each dNTP upon which reverse transcriptase (Promega; 40 units) was added. cDNA synthesis was allowed to proceed for 10 min at 37°C.

#### pipB2

To an annealing mixture (10 mM Tris-HCl (pH 8.0), 1 mM DTT, 60 mM KCl, 10 mM MgCl2 in DEPC-treated water), 0.05 pmol of in vitro-synthesized *pipB2* RNA was added, without or with increasing concentrations of RNA oligos EsvBr or EsvBg (IDT) and 0.6 pmol of end-labeled *pipB2*-specific primer (4054) were incubated for 10 min at 70°C. thereafter the tubes were transferred to RT for 15 min. The mixtures were then further incubated incubated for another 10 min at RT in 20 mM Tris–HCl, 10 mM magnesium acetate, 0.1 M NH_4_Cl, 0.5 mM EDTA, 2.5 mM β-mercaptoethanol (Sigma, cat. no. 8057400250), and 0.5 mM each dNTP upon which reverse transcriptase (Promega cat. no. M170A; 40 units) was added. cDNA synthesis was allowed to proceed for 10 min at 37°C.

The extension products were separated on 6% acrylamide 8 M urea-sequencing gels alongside sequencing reactions carried out with the same end-labeled primers (4054/4280) on pGEM-*siiA*/*pipB2* plasmids.

### β-galactosidase assays

Overnight cultures, as indicated, were diluted 1:100 and grown to the indicated growth condition in LB media containing appropriate antibiotics and 0.4-1 mM IPTG to induce expression from P*lacO* promoter. β-galactosidase activity was assayed as described (Basu *et al*.).

### STM0327 and EsvA stability assay

To estimate the half-life of *EsvA* and STM0327 mRNA, overnight cultures of SL1344 lacI^q^ carrying P*lac-*EsvA plasmid, as indicated, were diluted 1:100 into 70ml LB media in 250ml flasks supplemented with appropriate antibiotics and incubated at 37°C with shaking (200 rpm). After 80 min, 0.4 mM IPTG was added and after 45 min the cultures were split into two 35 ml subcultures, and rifampin (0.2 mg/ml) was added to one of them. Samples were taken for RNA extraction, before rifampin was added (0 min), and after the addition at 1, 5, 10, 20 min. RNA extraction was carried out using TRI reagent (Sigma, cat. no. T9424), and 5 µg of total RNA were run on polyacrylamide gel as described in Northern blot analysis section. Primer 4257 was used for hybridization.

### In vivo *pipB2* primer extension

Total RNA (30 µg) extracted using TRI reagent (Sigma, cat. no. T9424) from strains as indicated was incubated with an end-labeled *pipB2*-specific primer (3957/4116) at 70°C for 5 min, followed by 10 min on ice. The reactions were subjected to primer extension at 42°C for 45 min using 1 unit of MMLV-RT (Promega, cat. no. M170A) and 0.5 mM of dNTPs. Extension products were analyzed on 6% acrylamide 8 M urea-sequencing gels next to sequencing reactions primed with the same end-labeled primer carried on pGEM3-*pipB2*.

### SiiE secretion

Cultures of SL1344 lacI^q^ *siiE*::HA carrying P*lac* or P*lac-*EsvA plasmid or SL1344 lacI^q^ carrying P*lac* plasmid, as indicated, were grown overnight (18 h) in 10 ml LB medium supplemented with ampicillin at 25°C without agitation. Cultures were then diluted 1:10 into 30 ml of fresh LB medium in 125 ml flasks containing ampicillin and induced with 0.4 mM IPTG. The cultures were incubated at 37°C with shaking (200 rpm). OD_600_ was measured at 1, 2, and 4 hours post-induction to normalize protein concentrations. At the indicated time points 0.5–1 ml aliquot was collected and centrifuged for 5 minutes at maximum speed at 4°C. The pellets were resuspended in a calculated volume of 1X LDS sample buffer (GenScript, cat. no. M00676), supplemented with 2.5% β-mercaptoethanol (Sigma, cat. no. 8057400250), heated at 95°C for 5 minutes, and stored at -20°C until further use. In-parallel to isolate secreted proteins, the remaining culture volumes were transferred to chilled 50 ml tubes and centrifuged at 5,000 rpm for 10 minutes at 4°C. The supernatants were collected and passed through a 0.45 µm filter to ensure the removal of residual intact cells. Proteins were precipitated by adding 10% (v/v) trichloroacetic acid (TCA) (Sigma, cat. no. T6399), followed by vigorous vortex and overnight incubation on ice. Precipitated proteins were recovered by centrifugation at 12,000 rpm for 15 minutes at 4°C. The protein pellets were washed with 1 ml of cold acetone. Following a subsequent centrifugation at 5,000 rpm for 10 minutes at 4°C, the acetone was removed and the pellets were air-dried for 5 minutes. The dried pellets were resuspended in 30 µl of 1X LDS sample buffer supplemented with 2.5% β-mercaptoethanol. heated at 95°C for 5 minutes. For analysis, 5 µl of each sample were loaded onto gels for SDS-PAGE and ran for 3 hours.

### Total protein extraction

For SL1344 lacI^q^ *orgB*-SPA carrying P*lac* or P*lac*-EsvA plasmids, overnight cultures were diluted 1:100 into completely filled 15 ml tubes containing fresh LB media supplemented with ampicillin and 0.4mM IPTG and incubated at 37°C without shaking. After 3 hours cultures were removed to 125 ml flacks and incubated at 37°C with shaking (200 rpm) for 30 minutes. For SL1344 lacI^q^ *sicA*-SPA carrying P*lac* or P*lac*-EsvB plasmids, overnight cultures were diluted 1:100 into 10ml LB media in 125 ml flasks supplemented with ampicillin and incubated at 37°C with shaking (200 rpm). After 4 hours cultures were centrifuged at room-temperature and resuspended with fresh LB pH 5 media (adjusted with HCl) supplemented with 0.4 mM of IPTG, and let grow for additional 6 hours. At indicated time point, cultures were centrifuged for 5 minutes at 4°C and pellets were resuspended with 1X laemmeli buffer supplemented with 5% β-mercaptoethanol. Samples were then heated at 95°C for 10 minutes. laemmeli buffer volume for each sample were normalized to OD_600_.

### Western blot and Immunodetection

Following electrophoresis, proteins were transferred to a nitrocellulose membrane using the GenScript protein transfer system with standard protocol. After transfer, the membrane was incubated in a blocking solution (4% BSA and 4% skim milk in 1X TBST) for 1 hour at room temperature with constant shaking. The membrane was rinsed with 1X TBST before the addition of the primary antibody. For SiiE::HA detection, rabbit anti-HA antibodies were used. For OrgB-SPA, SicA-SPA and PipB2-SPA detection, mouse anti-FLAG antibodies were used. Membrane was incubated with primary antibody (diluted 1:5000 in 1X TBST) for 1 hour at room temperature. The membrane was subsequently subjected to three 10-minute washes with 1X TBST. Following the washes, the membrane was incubated with an HRP-conjugated secondary antibody (diluted 1:10,000 in 10 ml of 1X TBST) for 1 hour at room temperature with shaking. Washing was then repeated. Protein bands were visualized using Enhanced Chemiluminescence (ECL). Equal volumes (3 ml each) of ECL reagents (Advansta, cat. no. K-12045-D50) were mixed and applied to the membrane in the dark, followed by a 1-minute incubation with gentle shaking. The membrane was then blotted dry and subjected to automated imaging for signal exposure. Band normalized intensity was calculated using the Image Lab software (Version 6.1, Bio-Rad Laboratories).

### Adhesion assay

Overnight cultures of SL1344 lacI^q^ and SL1344 lacI^q^ ΔSTM0327::cm carrying P*lac* or P*lac-esvA* plasmids, were diluted 1:36 into 18ml glass tubes containing 3.5ml LB media, supplemented with appropriate antibiotics and 0.4 mM IPTG. The cultures were grown at 37°C on a rotor drum (60 rpm) for 3.5 hours upon which, cultures were diluted 1:10 in fresh LB medium. OD_600_ was measured and the required volume of bacterial suspension was used to infect HeLa cells or MDCK cells (MOI 1:5) and for plating on LB plates (to recalculate accurate CFU used for infection). Infected cultures were incubated at 37°C for 20 to 25 minutes to facilitate adhesion. Following incubation, wells were washed three times with 1 ml warm 1X PBS, and harvesting was done by adding 0.5 ml of PBS containing 1% Triton X-100. The lysate was pipetted up and down several times to ensure complete cell lysis, transferred to glass tubes and vortexed for 10 seconds, followed by incubation at room temperature for 10 minutes. Lysate was serially diluted with 1X PBS and appropriate amounts were plated on LB plates. CFU was counted and adhesion was calculated.

### SopD2 intracellular secretion assay

Overnight cultures of SL1344 lacI^q^ and SL1344 lacI^q^ Δ*pipB2*::FRT carrying P*BAD-sopD2-FLAG-HiBit* and P*lac* or P*BAD* or P*BAD-pipB2* plasmids, were diluted 1:100 into 10ml LB media in 125ml flasks supplemented with appropriate antibiotics and incubated at 37°C with shaking (200 rpm) until OD_600_ = 2. Cultures were diluted 1:100 in fresh LB medium and the number of bacteria required was calculated according to a MOI of 1:10. The appropriate volume of bacteria was added to HeLa LgBit seeded in parallel 24-well (Thermo Scientific, cat. no. 142475) and white 96-well plates (Thermo Scientific, cat. no. 136101). Plates were incubated for 30 min at 37°C in a 5% CO₂ to allow infection. Wells were washed twice with 1 ml warm 1×PBS, and fresh media was added (DMEM supplemented with 10% FCS and 100 µg/ml gentamicin; 100 µl per well for 96-well plates or 1 ml per well for 24-well plates). After 45 minutes, wells were washed twice with warm 1×PBS and replaced with no phenol-red DMEM (Sigma, cat. no. D8537) supplemented with 10% FCS, 1 mM IPTG, 10 µg/ml gentamicin, and drkBit (Promega, cat. no. N7400). 5 hours post-infection, 0.2% arabinose and Nano-Glo® Live Cell Reagent (Promega, cat. Number N2011) were added to the 96-wells plates. Plates were transferred to a plate reader, and luminescence was measured every 15 min. At the same time, the parallel 24-well plates were washed twice with 1X PBS and lysed with 1% triton X-100. Bacteria cells were serially diluted with 1xPBS and were plated for CFU enumeration. RLU was normalized to CFU.

### SopD2 secretion, pH 5 assay

Overnight cultures of SL1344 lacI^q^ and SL1344 lacI^q^ Δ*pipB2*::FRT carrying P*lac* or P*lac-*EsvB, were diluted 1:100 into 10ml LB media in 125ml flasks supplemented with appropriate antibiotics and incubated at 37°C with shaking (200 rpm). After 4 hours, cultures were centrifuged at 5000rpm for 5 minutes at room-temperature and resuspended with 10ml of fresh LB media adjusted to pH 5 with HCl as described (Gong *et al*.). After additional 5.5h of growth, 0.2% arabinose was added, and cultures were grown for additional 30 minutes. OD_600_ was measured and 25µl of each culture was transferred to a white 96-well plate (Thermo Scientific, cat. no. 136101). An equal volume (25µl) of buffer mix (buffer containing substrate and LgBit, Promega, cat. no. N2420) was added to each well, plates were briefly shaken, and luminescence was measured. RLU was normalized to OD_600_.

